# A Phylogeny for Heterostraci (stem-gnathostomes)

**DOI:** 10.1101/2022.08.11.503478

**Authors:** Emma Randle, Joseph N. Keating, Robert S. Sansom

## Abstract

The armoured jawless fishes (ostracoderms) are major and widespread components of middle Palaeozoic ecosystems. As successive plesia on the gnathostome lineage, they reveal the early sequences of vertebrate evolution, including the assembly of the vertebrate skeleton. This is predicated however, on understanding of their diversity and interrelationships. The largest ostracoderm clade, the Pteraspidimorphi, is often reconstructed as sister taxon to other boney vertebrates yet they lack a phylogenetic framework, in particular the heterostracans. Problematic heterostracans with a tessellate headshield (‘tessellate-basal’ model) are often regarded as the plesiomorphic condition for the clade but no phylogenetic analysis has included these taxa. Here we review the Heterostraci and present their first comprehensive phylogenetic analysis (131 heterostracan taxa and 12 outgroup taxa). Heterostraci and Ordovician Pteraspidimorphi are recovered as sister-group to all other boney jawless vertebrates in parsimony analyses, however, in no instances do we recover a monophyletic Pteraspidimorphi. Tree visualization reveals lack of resolution results from two conflicting solutions for the heterostracan ‘root’. Stratigraphic congruences provides support for the macromeric Ctenaspisdidae as sister taxon to all other Heterostraci rather than the “tesselate-basal” model. The results presented here are the first phylogenetic hypotheses of heterostracan relationships and it is hoped a first step into an accurate interpretation of character evolution and polarity in this crucial episode of vertebrate evolution.

## 1. INTRODUCTION

Relationships of early vertebrates are important to understanding the early events in our own evolutionary history. The armoured jawless vertebrates known as the ‘ostracoderms’ are key in the context as they bridge the gap between living jawless vertebrates i.e. hagfish and lamprey, and the jawed vertebrates (gnathostomes) (Fig. 1). Ostracoderms span an important part of the gnathostome stem lineage and reveal the acquisition of many definitive gnathostome features including paired appendages, bone and nasal organs (Donoghue & Keating, 2014). The phylogenetic context of these taxa are therefore extremely important to understanding the sequence and timing of vertebrate innovations. Many ostracoderm taxa have a phylogenetic framework in place. For example, osteostracans are resolved as the sister group to jawed vertebrates and their internal relationships indicate the non-cornuate ateleaspids are the deepest branching osteostracans (Sansom, 2009a,b). Other ostracoderm groups such as the Thelodonti (Wilson & Märss, 2009), Anaspida (Blom, 2012; Keating & Donoghue, 2016), Galeaspida (Zhu & Gai, 2007, Gai et al., 2018) have phylogenetic trees. However, the taxonomically largest group of ostracoderms i.e. the Heterostraci still lacks a whole group analysis. This serves as a major barrier to understanding not only their diversity through time and space, but also has ramifications for hypotheses of vertebrate evolution.

**Fig. 1.**
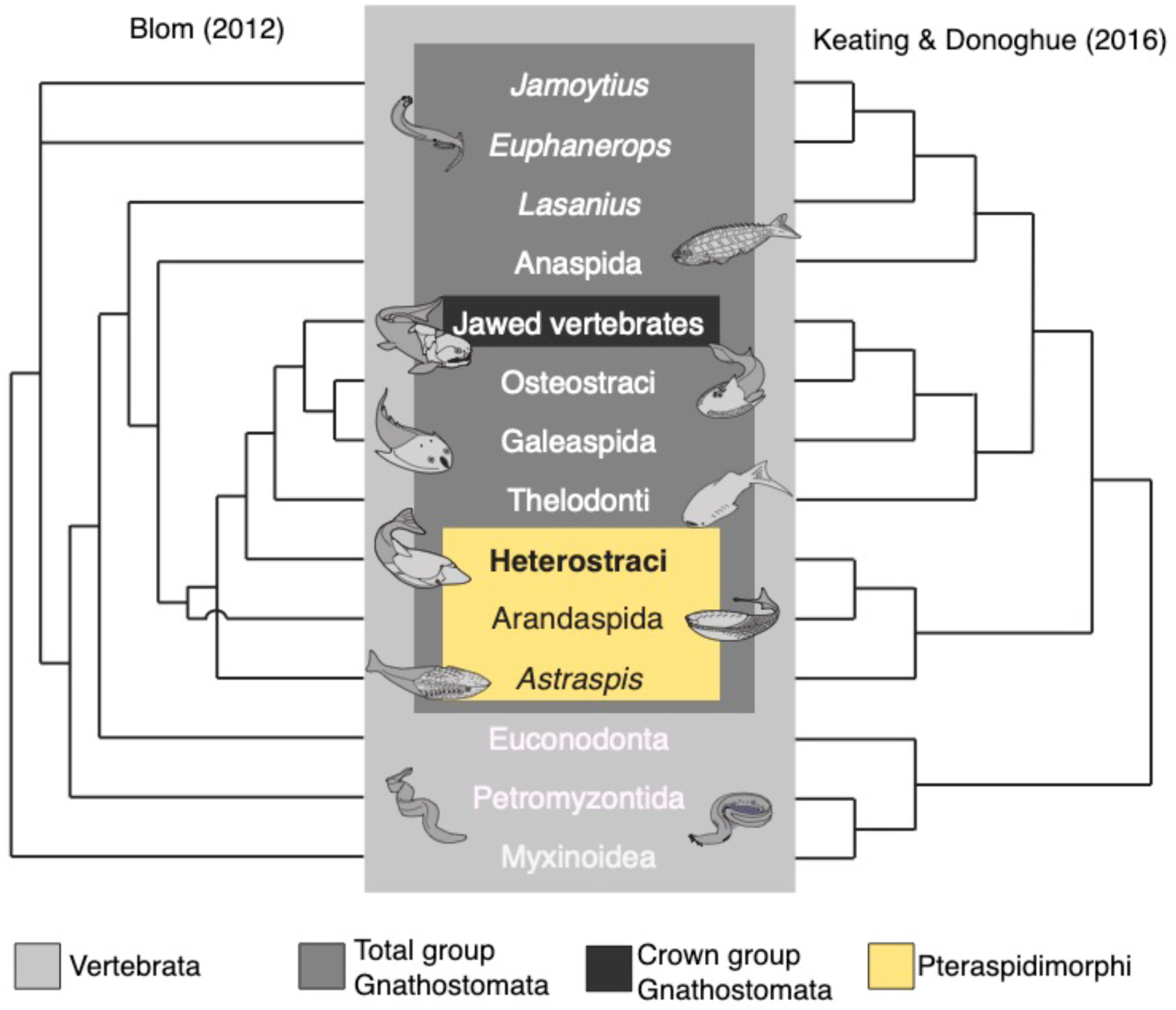
Conflicting vertebrate phylogenies of Blom (2012) and Keating & Donoghue (2016).

Heterostraci dominated vertebrate assemblages from the Middle Silurian to the Early Devonian (Purnell, 2001; Anderson *et al*., 2011; Sansom, Randle, & Donoghue, 2015) before their decline and extinction toward the end of the Devonian (Randle, & Sansom, 2019). Their position on the gnathostome-stem is still debated, with recent phylogenies providing conflicting evidence (Fig. 1). Heterostraci are generally reconstructed as one of the earliest branching stem-ganthostome clades as part of the Pteraspidimorphi (i.e. sister taxon to jawed vertebrates + osteostracans + galaeaspids + thelodonts, Pteraspidimorphi as either monophyletic or paraphyletic). As such, they are important for scenarios of skeleton evolution (Blom 2012, Keating & Donoghue, 2016, Miyashita et al., 2019, 2021, Sansom et al., 2010). There is however, conflict over the root of gnathostomes, and the position of heterostracans relative to anaspids and other Pteraspidimorphi.

This conflict in position has widespread ramifications for interpreting the ancestral vertebrate condition. A possible reason the position of heterostracans on the vertebrate stem is ambiguous is because they lack a broad encompassing taxonomic phylogeny. Many key groups such as the tessellate heterostracans, Amphiaspididae and Traquairaspididae have yet to be subjected to a cladistic analysis, with previous analyses focusing on smaller clades within the Heterostraci i.e. Pteraspidiformes and Cyathaspididae (Ilyes & Elliott, 1994; Pernègre, 2002; Pernègre & Goujet, 2007; Pernègre & Elliott, 2008; Lundgren & Blom, 2013; Randle & Sansom, 2016, 2017) or Psammosteoidei (Glinskiy 2017).

Here we undertake the first comprehensive phylogeny for heterostracans with the aim to not only resolve the intra-relationships of the Heterostraci, but also their position with the gnathostome stem. Our taxon sample therefore is a comprehensive selection of heterostracan genera (n=131) and a wide range of outgroup taxa i.e. Ordovician pteraspidimorphs (*Sacabambaspis*, *Astraspis*, *Arandaspis*) and other ostracoderm taxa. To support our analyses, we survey heterostracan anatomy and diversity, and previous interpretations of relationships. We construct a comprehensive list of relevant characters for parsimony and Bayesian analysis, before we finally consider the implications of the phylogeny for heterostracan evolution, and vertebrate evolution.

## 2. HETEROSTRACAN ANATOMY

The bone of heterostracan dermal skeleton is composed mainly of aspidin, a type of acellular bone (Keating et al 2018), which covered their head and tail regions. Their headshields were encased in large plates (sometime composed of smaller boney platelets or tesserae), which would have projected their internal anatomy. The trunk region was covered in rhombic or elongate boney scales. Heterostracans lacked paired appendages or fins, and as such the caudal fin was their only means of muscular propulsion. The Heterostraci have previously been defined as a clade based upon possessing a single pair of branchial openings, having a dermal skeleton composed of aspidin, which is often but not always topped with dentine and enameloid (Keating, Marquart, & Donoghue, 2015). Here we survey heterostracan anatomy in order to support the construction of a character list for phylogenetic analysis. This allows clarification of terminology and interpretations of homology.

### Dorsal plate & shield

Heterostracans, like other stem-gnathostomes, are distinguished by having an ossified headshield, which is posteriorly delimited by the body and caudal regions (usually covered in elongate or diamond shaped scales). A distinction is made here between the dorsal plate (Fig. 2) and dorsal shield (Fig. 3). The dorsal shield is a singular plate that encompasses the pineal organ, covers dorsally the rostral region, and delimits fully or partially the orbital area (for those with an external orbital opening), whereas the dorsal plate is a separate plate that does not cover these regions (for which there are separate plates). In some, the dorsal shield extends laterally forming a lateral brim (Fig. 3D)(covered dorsally and ventrally by ornament). In cyathaspids the dorsal shield folds anteriorly to form the maxillary brim, and the anterior margin of the orbital opening (Fig. 3A,C). Laterally the shield forms the posterior border of the orbital opening and branchial openings, with the post-branchial lobe delimiting the posterior edge of the branchial opening. In other forms the dorsal shield delimits completely the orbital opening and branchial openings e.g. *Natlaspis* (Fig.3G). The headshield of tessellate heterostracans is regarded here as homologous to the singular headshield seen in cyathaspids. In the tessellate heterostracan *Lepidaspis* (Fig. 3F), the tesserae have fused into a single headshield that covers the pineal, rostral and orbital areas.

**Figure 2.**
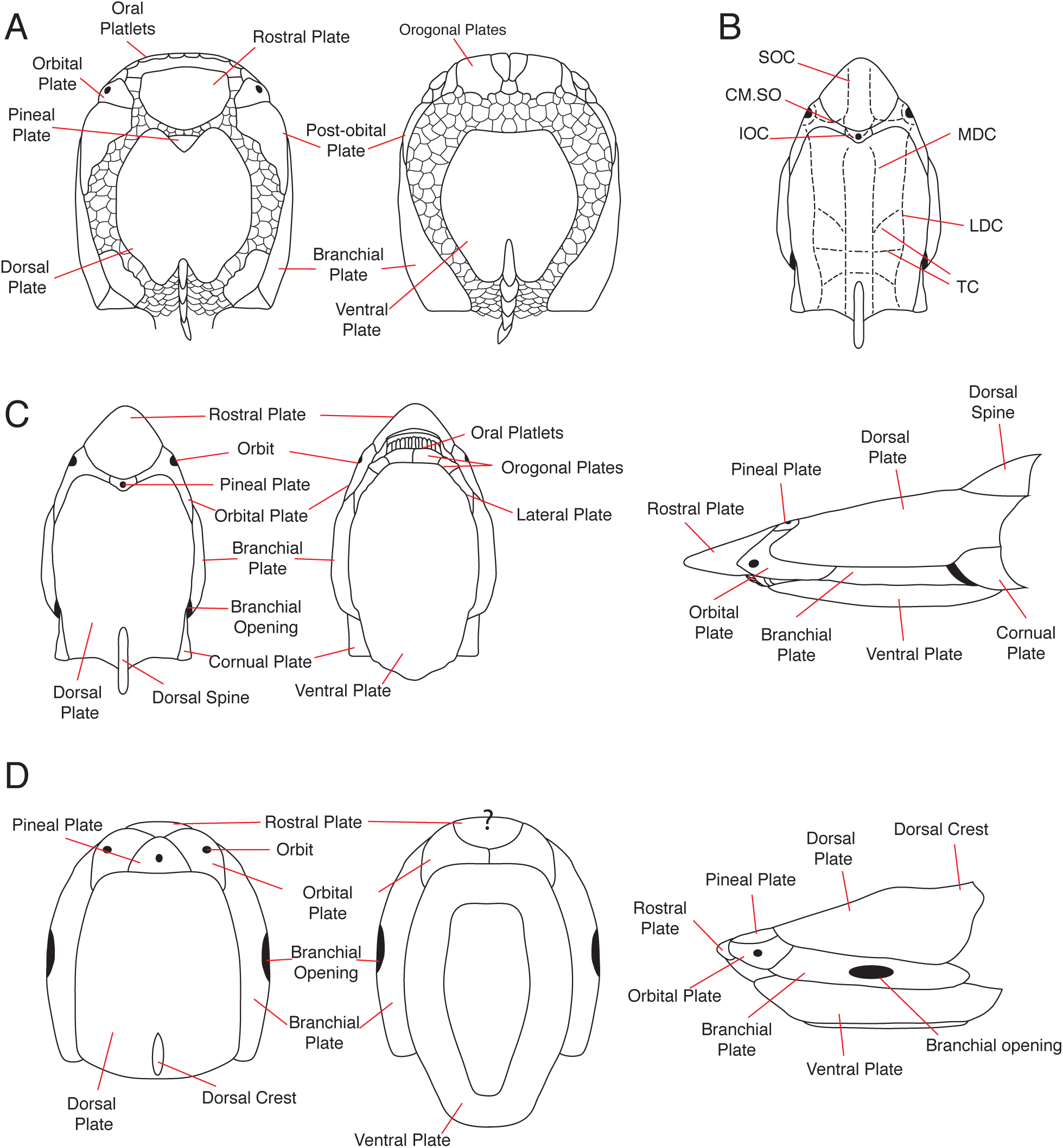
Cartoons of heterostracan anatomy in taxa that have separate plates. A *Drepanaspis* psammosteid adapted from Obruchev & Mark-kurik (1965). B sensory canal pattern: SOC, supraorbital canal; CM.SO, supraorbital commissure; IOC, inter-orbital canal; LDC, lateral dorsal canal; MDC, medial dorsal canal; TC, transverse commissures of a pteraspid adapted from Randle & Sansom (2016). C cartoon of a pteraspid adapted from Randle & Sansom (2016). D. *Toombsaspis* a traquairaspid adapted from Tarrant (1991).

**Figure 3.**
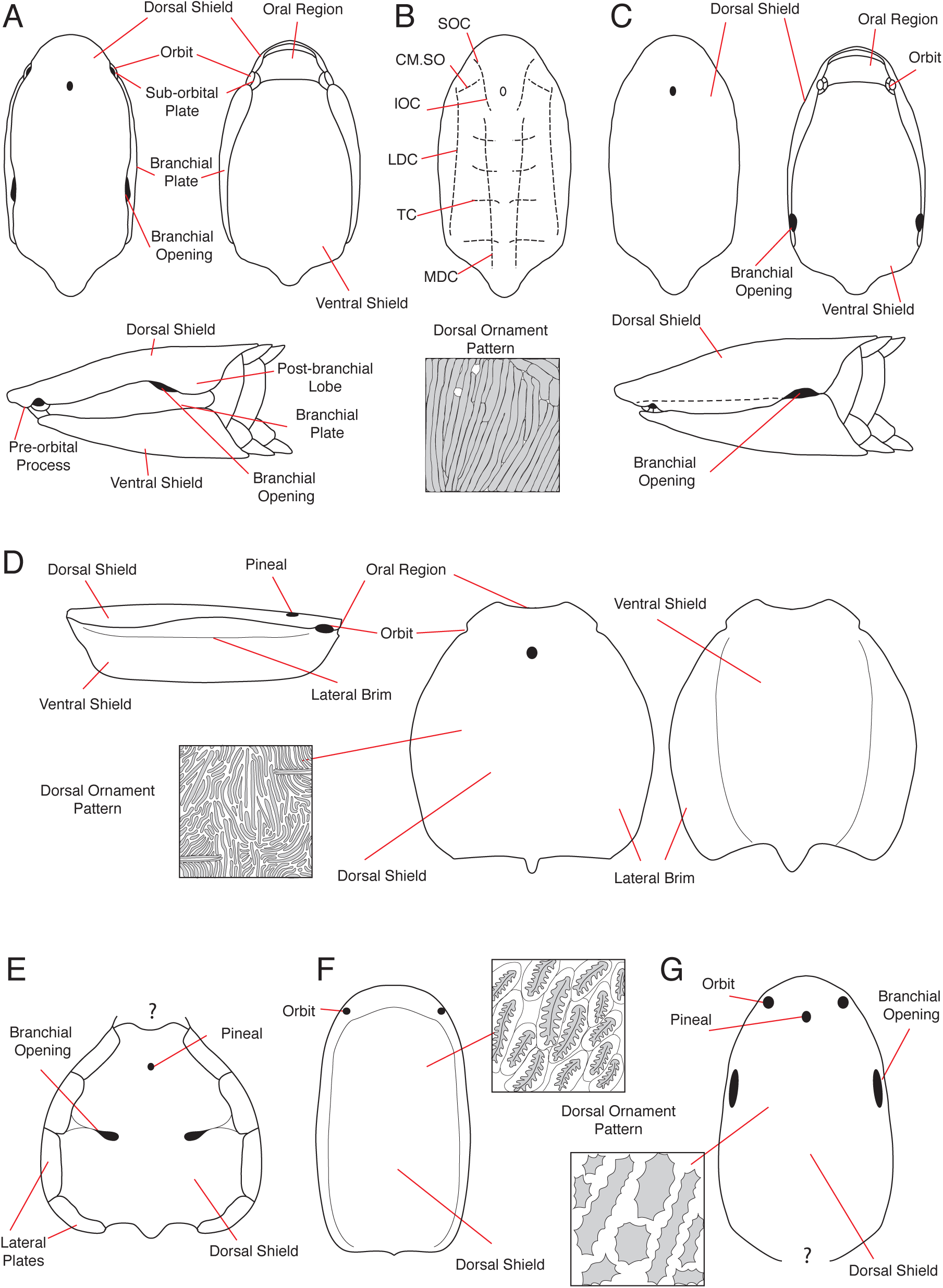
Cartoons of heterostracans with a dorsal shield. A poraspid-like cyathaspid adapted from Kiaer (1932), Denison (1964) and Randle & Sansom (2017). B general cyathaspid sensory-line canal pattern: SOC, supraorbital canal; CM.SO, supraorbital commissure; IOC, inter-orbital canal; LDC, lateral dorsal canal; MDC, medial dorsal canal; TC, transverse commissures from Randle & Sansom (2017). C *Allocryptaspis*-like cyathaspid adapted from Denison (1964) and Randle & Sansom (2017). D *Siberiaspis* amphiaspid. E. *Cardipeltis* a problematic heterostracan. F. *Lepidaspis* a tessellate heterostracan. G. *Natlaspis* a problematic heterostracan.

### Pineal Organ

The pineal organ is regarded as a light sensitive spot and can usually be found at the anterior end on the dorsal side of the headshield. In some taxa (Pteraspidiformes, Psammosteidae and Traquairaspididae) the pineal organ is encompassed in the pineal plate (Fig. 2), whereas in others it is encased within the dorsal shield (Fig. 3). The pineal organ is often covered by a layer of superficial dentine and enameloid, and not undercut by dermal bone (Keating *et al*., 2015). The pineal is not apparent in all taxa.

### Rostral area

The rostral region extends anteriorly from the orbital openings. It can be extensive in some taxa (e.g. *Rhinopteraspis*) and non-existent in others (e.g. *Siberiaspis*). In Pteraspidiformes and Psammosteidae, the rostral area is covered by a distinct separate plate. In Pteraspidiformes the rostral plate folds over laterally and anteriorly to form the ventral side of the plate. Seen on the ventral side of the rostral plate in Pteraspidiformes are the pre-oral surface and pre-oral fields. The rostral plate in Psammosteidae is anteriorly delimited by the dorsally-orientated oral region, it is therefore only ornamented on the dorsal surface. In forms with a dorsal shield, the rostral area of the shield can be lacking or truncated (seen in amphiaspids such as *Kureyaspis*), or can fold anteriorly to form the ventrally orientated maxillary brim (analogous to the pre-oral surface of forms with a rostral plate). In some unusual amphiaspid forms the rostral and oral region extend anteriorly to form a ‘tubular mouth’ (e.g. *Eglonaspis*).

### Orbital area

The majority of heterostracan taxa have a pair of orbital openings, which are surrounded by an orbital plate in Pteraspidiformes, Psammosteidae and Traquairaspididae (Fig. 2). In forms with a dorsal shield the orbits are enclosed by the dorsal shield, as seen in Amphiaspididae and problematic taxa (Fig. 3D-G), or partially delimited by the dorsal shield in the Cyathaspididae Fig. 3A-C).

### Branchial openings

Having a single pair of branchial openings is a supposed synapomorphy of the Heterostraci. A number of different clades possess a branchial plate, which when present precedes the branchial opening. The plate is found in the Pteraspidiformes, Psammosteidae, Cyathaspididae and Traquairaspididae (Fig. 2, 3A), and is treated here as a homologous feature. Not all taxa have a branchial plate. There are many different combinations of plates that delimit the branchial opening. However, if a taxon has a branchial plate, then it inherently will delimit the branchial opening.

### Oral region

The oral region for many heterostracans is not preserved. For those taxa that it is preserved, there is a complex array of plates that do not provide much information regarding the feeding habits of these jawless boney vertebrates (although see Purnell 2002). For articulated specimens without oral plates such as the Amphiaspididae, it is unclear whether these plates were absent, have not preserved, or were a non-boney element. In the phylogenetic analyses presented here, the larger oral plates found in heterostracans are named orogonal plates and the finger-like smaller plates are regarded as the oral platelets. However, the oral platelets of Psammosteids (e.g. *Drepanaspis*) are larger and wider than those seen in Pteraspidiformes but still possess the characteristic hook shape, therefore are considered oral platelets (Fig. 2A) (Gross, 1963).

### Body Region

The body and caudal region are rarely preserved articulated with headshields for heterostracan taxa. The dorsal and ventral margins have larger ridge scales (sometimes referred to as fulcral scales), which are more curved and elongate than the lateral scales. Some taxa have a single row of these, whereas other taxa have up to three rows (*Dinaspidella*). Body scales can be small and diamond shaped or more elongate and rectangular (*Nahanniaspis, Allocryptaspis*). The shape of the caudal fin has long been debated in the heterostracan literature. Generally, the caudal tail of heterostracans’ appears to be equi-lobed or with a more pronounced ventral lobe, with thicker radials (scales) surrounded by much finer scales (Denison, 1971; Soehn & Wilson, 1990; Pellerin & Wilson, 1995; Mark-Kurik & Botella, 2009).

### Sensory Canals

The sensory canal system in heterostracans is most commonly found in the reticulate layer (sitting just below the ornament layer) within the dermal armour. It can often be seen on the external ornamented surface as a network of pores (as seen in Cyathaspididae and Pteraspidiformes) or grooves (as seen in Amphiaspididae). Some heterostracans, namely Amphiaspididae (and potentially some Psammosteidae and Traquairaspididae) have a second deeper canal system found in the spongy bone layer and with a different pattern to the sensory canal system found in the reticulate layer (Novitskaya, 1983; Elliott & Mark-Kurik, 2005). Sensory canals of the Heterostraci are often used in taxonomic distinction. Some differences between major clades include; the two arms of the pineal/inter-orbital canals, in Cyathaspididae do not meet to form a singular canal as they do in the Pteraspidiformes, the Pteraspidiformes have 3 pairs of radially arranged transverse commissures, whereas, other clades have 4 pairs arranged in parallel (Fig. 2B, 3B).

### Histology

Heterostracans’ vertebrate affinity was first recognised after Huxley (1858) undertook a histological examination of their dermal skeletons, which preserve a unique micro-texture. Heterostracans lack osteocytes (bone cell spaces) and instead their bone is composed of a type of acellular bone named aspidin (Keating *et al*., 2015, Keating et al 2018). Histological examination of heterostracan dermal skeletons using X-ray synchrotron tomography identified a 4 layer structure (Keating *et al*., 2015) comprising a superficial ornament layer (orthodentine and enameloid) topping the 3 dermal bone layers including a reticulate, middle spongy layer and a dense (plywood-like) basal layer.

### Ornament

Different patterns and properties are seen in the superficial ornament layer of heterostracans. The ornamented part of the dermal shield is composed of dentine capped with enameloid (Keating *et al*., 2015), except in the ctenaspids and obruchevids, which appear to lack any superficial dentine (Elliott et al. 2004; Elliott & Blieck 2010). Ornament varies from continuous uniform ridges seen in Pteraspidiformes and Cyathaspididae to stellate ornament of Psammosteidae and “oak leaf” shaped ornament seen in *Lepidaspis*. Tops of tubercule ornament vary from peaked, ridged or flat-topped and have smooth, crenulated or scalloped edges. The density of ornament can differ from very fine densely packed ridges (20/mm) to large singular well-spaced tubercules (each>1mm) separated with large gaps of dermal skeleton barren of ornament. Areas demarked by changes in the superficial ornament pattern (coined epitega) are see in some cythaspids (Denison, 1964). These areas possible foreshadow the separate plates of other heterostracans and likely demark different growth centres. The epitega are sometimes marked by a suture or a band of smaller rounded dentine tubercules (Denison, 1964) and the different regions can have different relief to those in adjacent areas. As ornament is used in taxonomic distinctions, an attempt to characterize this variation is given below. It remains to be seen *post hoc* if this is a phylogenetically informative character or if it is homoplastic. It is most likely useful for distinguishing taxa at the species level once other characters/features have been used to determine taxonomic affinity.

## 3. HETEROSTRACAN DIVERSITY

Within the Heterostraci are a number of smaller clades predominantly at order or family level. These include: Pteraspidiformes, Cyathaspididae, Psammosteidae, Amphiaspididae, and Traquairaspididae. There are also a number of heterostracan taxa of uncertain affinity (problematica). Below we will describe and detail the main classes of heterostracan anatomy used in the phylogenetic analysis. For each we consider i) their composition and defining features, ii) previous phylogenetic interpretations, and iii) a brief description of stratigraphic range and palaeoenvironment. We also consider the non-heterostracan pteraspidimorphs as they are important out-group taxa for our analyses.

### Pteraspidiformes

The Pteraspidiformes are taxonomically the largest clade and arguably the most iconic group of heterostracans, having received the most attention with regards to their intra-relationships (Ilyes & Elliott, 1994; Pernègre, 2002; Pernègre & Goujet, 2007; Pernègre & Elliott, 2008; Randle & Sansom, 2016). There are approximately 46 named genera and over 110 species (Randle & Sansom, 2016). Characteristics of the Pteraspidiformes include: separate dorsal, ventral, rostral and pineal plates, along with paired orbital, branchial and often cornual plates (Fig. 2B-C) (Blieck, 1984) all of which fuse on maturity (Denison, 1960; Blieck *et al*., 1991; Elliott, Schultze, & Blieck, 2015). Regions useful in taxonomic identification include the shape of the pineal plate, pineal-orbital belt, shape of rostral plate, pattern of the sensory canals, and the shape and proportions of the median dorsal plate (Elliott, 1983, 1984; Blieck, 1984; Blieck *et al*., 1991).

The anchipteraspidid Pteraspidiformes were believed to be a link between Cyathaspididae and the Pteraspidiformes, and do indeed share features that appear to transition between the two groups including a pineal plate enclosed within the dorsal plate/shield, a closed inter-orbital canal that loops around the pineal organ and concentric ornament patterns (Elliott, 1984). However, in the phylogenetic analysis of (Randle & Sansom, 2017) the Anchipteraspididae were placed as sister group to the Pteraspidiformes when coded as pteraspid-like. When they were coded as cyathaspid-like, their pteraspid affinity became ambiguous as they fell in a large polytomy with many other cyathaspid taxa.

An early distinction within the Pteraspidiformes was between the Protopteraspididae and the Pteraspididae (Blieck, 1984). The Protopteraspididae contained some of the oldest pteraspids including *Protopteraspis, Stegobranchiaspis* and *Doryaspis* and was generally defined as having an inter-orbital canal that loops around the pineal plate. The Pteraspididae were seen as more derived and had an inter-orbital canal that looped through the pineal plate. Pernègre & Elliott (2008) subdivided the Pteraspidiformes up even further after their phylogenetic analysis identified two further clades within the Pteraspididae of Blieck (1984), namely the Protaspididae – containing wide shielded often tuberculated pteraspids mainly from the Western-USA and a more limited Pteraspididae containing taxa not belonging within this wide shielded clade. However, previous phylogenetic analyses of Pteraspidiformes have not included all Pteraspidiformes – but only those with articulated or well know anatomies. Randle & Sansom (2016) included all taxonomically described Pteraspidiformes and were the first to utilize quantitative characters to reconstruct their phylogenetic relationships. They recovered a number of families including the Protopteraspididae, Protaspididae and Rhinopteraspididae. However, the phylogenetic affinity of many Pteraspidiformes taxa remained uncertain. Also demonstrated to belong within Pteraspidiformes are the Psammosteidae. Indeed when included within phylogenetic analyses of Pteraspidiformes they have been found positioned between the Anchipteraspididae and non-Anchipteraspididae Pteraspidiformes (Pernègre & Elliott, 2008), in a highly derived position within the Pteraspidiformes (Pernègre, 2002, Glinskiy 2017) or nested somewhere in between (Randle & Sansom, 2016; 2017).

The first fossil record occurrence of Pteraspidiformes is in the Prìdoli (Upper Silurian) of Arctic Canada. However, the Lochkovian (Lower Devonian) is when Pteraspidiformes appear to diversify and appear in high numbers. The youngest occurring Pteraspidiformes is *Helaspis* found in deposits of Givetian (Middle Devonian) age (Elliott, Dineley, & Johnson, 2000). Unlike other groups of heterostracans (i.e. Amphiaspididae), they do not show strong endemism and are found in a range of palaeocontinents including Laurentia, Avalonia, Baltica and Kara (Randle & Sansom, 2016). Pteraspidiformes are predominantly found in fluvial environments (Sallan et al. 2018).

### Cyathaspididae

Cyathaspididae are the second largest clade of heterostracans. Taxa traditionally included within the Cyathaspididae have a separate ventral and dorsal shield - the latter of which encompasses the pineal organ, rostral region and delimits partially the orbital opening (in some the orbit is underlain by a supraorbital plate). Some members have a pair of branchial plates, whereas in other taxa these are absent, and the branchial openings are delimited by the dorsal and ventral shields. The dorsal shield of some cyathaspids is superficially divided into discrete ornamented areas named epitega (Denison, 1964). Another characteristic feature of some cyathaspids is a lateral extension of the dorsal shield named the lateral brim and an ascending lamellae covering the lateral region (Denison, 1964).

The Cyathaspididae have historically been interpreted as monophyletic and derived members within the Heterostraci. Groups contained within Cyathaspididae include; Poraspidinae, Ctenaspidinae, Cyathaspidinae, Irregularaspidinae, Tolypelepidinae and Boothiaspididae, along with other taxa of uncertain affinity such as *Listraspis, Allocryptaspis* and *Ariaspis* (Denison, 1964; Elliott, Reed, & Loeffler, 2004; Elliott & Blieck, 2010; Elliott & Swift, 2010; Elliott, 2013, 2016). Janvier (1996) placed them as the sister group to the Amphiaspididae within the Cyathaspidiformes which in turn was the sister group to the Pteraspidiformes. Only a small number of phylogenetic analyses have been performed on the Cyathaspididae, with the first concentrated on their intra-relationships (Lundgren & Blom, 2013), and the second as part of an analysis containing Pteraspidiformes (Randle & Sansom, 2017). Randle & Sansom (2017) found the Cyathaspididae to be paraphyletic with respect to the Pteraspidiformes. However, no formal phylogenetic analysis to date contains the Amphiaspididae which are believed to be the sister group to the Cyathaspididae (Obruchev, 1967; Halstead, 1973; Novitskaya, 1983; Blieck *et al*., 1991; Janvier, 1996). Cyathaspididae taxa have an unusual combination of characters, which has led authors to align them with many different groups including Pteraspidiformes, Psammosteidae and Traquairaspididae (fig. 4). It is most likely the Cyathaspididae represent a generalist or plesiomorphic condition for the Heterostraci, yet this has not been tested.

**Figure 4.**
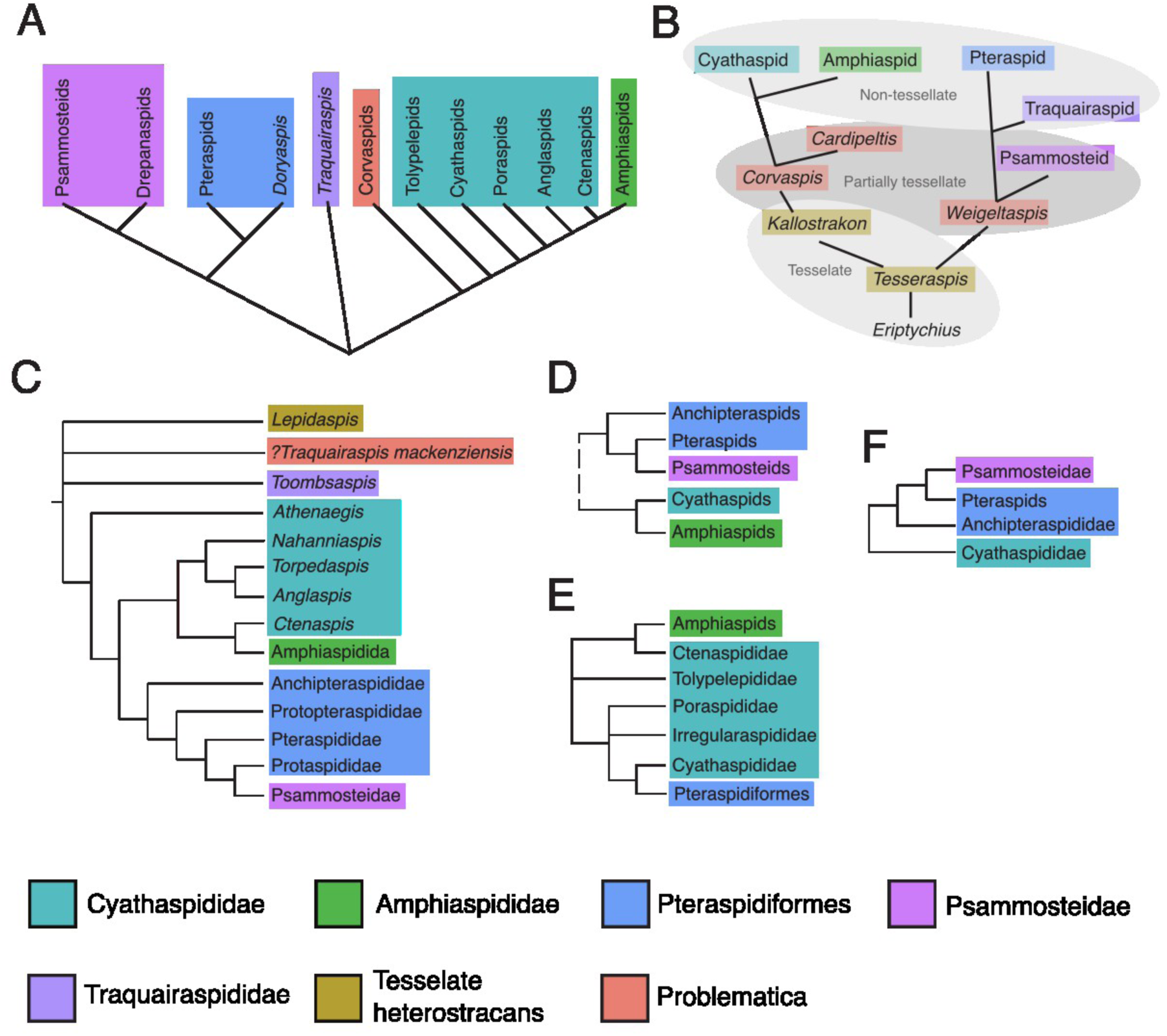
Previous hypotheses of Heterostraci intra-relationships. A. hypothesis of heterostracan relationships by Obruchev (1967). B. Halstead’s (1973) evolutionary trajectory of heterostracans from a tessellate ancestor. C. phylogeny as proposed byJanvier (1996). D. relationships envisioned by Blieck, Elliott, & Gagnier (1991). E. affinity of heterostracan taxa by Novitskaya (1983). F. Phylogenetic results of Randle & Sansom (2017) which included Cyathaspididae, Pteraspidiformes and some Psammosteidae.

Within the Cyathaspididae are some of the earliest members of the Heterostraci: *Athenaegis, Archaegonaspis* and *Tolypelepis* from the Wenlock (Middle Silurian) (Dineley & Loeffler, 1976; Märss, 1977; Soehn & Wilson, 1990). They are one of the dominant groups in heterostracan evolutionary history but are replaced by the Pteraspidiformes as the most abundant in the Lochkovian. Representatives of the Cyathaspididae survive until the Eifelian of Western USA. Similarly to the Pteraspidiformes they inhabit a number of palaeocontinents including Laurentia, Avalonia, Baltica and Kara.

### Psammosteidae

The Psammosteidae includes the stratigraphically youngest heterostracans and contains some of the physically largest members of the Heterostraci (*Obruchevia*, *Traquairosteus* and *Pycnosteus*). They are exemplified by *Drepanaspis* – the only known articulated psammosteid from the Hünsruck beds of Germany. *Drepanaspis* has been used to infer the arrangement of plates in other disarticulated psammosteids. Similar to the Pteraspidiformes, they have separate dorsal, ventral, pineal, branchial, orbital and rostral plates, along with post-orbital and adapted orogonal plates.

A number of Psammosteid families have been erected including the Drepanaspididae, Pycnostidae, Psammoleipididae, Psammosteidae and the Obrucheviidae (Tarlo, 1964, 1965; Elliott, Mark-Kurik, & Daeschler, 2004; Novitskaya, 2004). Tarlo (1964) considered the phylogenetic relationships of the psammosteids and envisioned a dichotomous split between the major groups, with the Pycnostidae in one group and the Psammolepididae, Psammosteidae and Obrucheviidae in another. A close affinity between the Psammosteidae and Pteraspidiformes has long been hypothesized. They both have the same basic configuration of plates; Gross (1963) demonstrated that juvenile *Drepanaspis* lack the superficial tesserae in between the larger plates which led many authors to believe that the Psammosteidae were derived from the Pteraspidiformes. This relationship is recovered in phylogenetic analyses that have included representatives of both clades (Pernègre, 2002; Pernègre & Elliott, 2008; Randle & Sansom, 2016, 2017). The hallmark in psammosteid relationships was achieved with the comprehensive analysis of Glinkskiy (2017). This analysis supported close relationships with the Pteraspidiformes and Drepanaspis as sister taxon to all other psammosteids.

*Drepanaspis* is the oldest Psammosteidae occurring in sediments of Emsian (lower Devonian) age (Gross, 1963; Tarlo, 1964). Psammosteids radiate and occur in the fossil record until the Frasnian (Upper Devonian) where they go extinct at the Frasnian/Famennian boundary. Psammosteids are found in Baltica, Laurentia and Kara palaeocontinents and can be found predominantly in fluvial environments (Gross, 1963; Tarlo, 1964; Obruchev & Mark-Kurik, 1965; Elliott *et al*., 2004).

### Amphiaspididae

The Amphiaspididae are an enigmatic group of heterostracans from the Northwestern Siberian Platform and the Taimyr Peninsula (Novitskaya, 2008). They are characterised by having singular dorsal and ventral headshields or a single headshield enclosing the whole of their headshield (Novitskaya, 1983, 2004; Janvier, 1996). None of the amphiaspid taxa possessed branchial plates, instead the branchial opening is encompassed within the dorsal shield or in forms in which no brachial opening have been found, most likely opened at the posterior of the headshield. Some amphiaspids have a post-orbital opening (pre-spiracular of Janvier (1996)) of unknown function, positioned laterally to the orbital openings (Novitskaya 1983). Another key characteristic is distinctive sensory grooves (rather than pores seen in other groups) that are sometimes delimited by ornament and often cut across the headshield and other ornament. Some amphiaspid taxa have a truncated rostral region and an anteriorly orientated mouth (e.g. *Siberiaspis*) and other forms have elongate or anteriorly extended rostral regions that are enclosed all around forming a tube-like mouth.

Amphiaspididae have traditionally been considered monophyletic and have a close relationship with the Cyathaspididae. However, they have yet to be included within any formal phylogenetic analysis (Obruchev, 1967; Halstead, 1973; Novitskaya, 1983; Blieck *et al*., 1991; Janvier, 1996). Similarities in gross morphology between *Liliaspis, Paraliliaspis* (both considered Cyathaspididae), and some Amphiaspididae taxa (i.e. *Putoranaspis*) perhaps imply a close relationship between the taxa. *Liliaspis* and *Paraliliaspis* (Cyathaspididae) both posses sensory pores and sensory grooves on their headshields that are characteristic of the Amphiaspididae. Amphiaspids also have the lateral brims found in some cyathaspids. However, amphiaspids differ in having their orbits and sometimes branchial openings enclosed in their dorsal shields. Novitskaya (2004) split the Amphiaspididae into 3 suborders, namely; Amphiaspidoidei which contains taxa possessing the post orbital opening, Hibernaspidoidei containing taxa with an elongate oral tube and Siberiaspidoidei which contains amphiaspid taxa which lack both the oral tube and post orbital openings.

The Amphiaspididae are only found in deposits from the Northwestern Siberian platform and Taimyr from the Lower Devonian (Lochkovian to Pragian) (Novitskaya, 1983, 2004, 2008). They inhabited marine to marginal marine environments, with Novitskaya (2008) attributing their inability to adapt to changing transgressive cycles as key to their demise.

### Traquairaspididae

The traquairaspids are more limited in diversity. They are characterized by having separate dorsal, ventral, pineal plates, and paired orbital and branchial plates, the latter of which are bisected by their branchial openings (Tarrant, 1991).

The Traquairaspididae have not previously been included in any phylogenetic analyses. Tarrant (1991) subdivided the traquairaspids into two families (Phialaspididae and Traquairaspididae), and placed them both in the order Traquairaspidiformes. When considering the evolutionary relationships of heterostracans, both Obruchev (1967) and Janvier (1996) (Fig. 4) placed the traquairaspids in an uncertain polytomy at the base of the Heterostraci, whereas, Halstead (1973) considered them to be fairly derived and the sister group to Pteraspidiformes (Fig. 4B). There are a number of problematic heterostracans from the Delorme Formation of Canada affiliated with the traquairaspids (i.e.*?Traquairaspis*) (Dineley & Loeffler, 1976). These Canadian problematica have a very different anatomy to Traquairaspididae (their dorsal headshield is composed of a singular plate encompassing both the orbital and branchial openings rather than separate plates). The species of *?Traquairaspis* most likely represent an as yet unnamed group of heterostracans.

Traquairaspids are found in sediments from Avalonia and Laurentia palaeocontinents. They have a similar stratigraphic distribution to cyathaspids, first appearing in the fossil record during the Wenlock (Middle Devonian) and surviving until the Lochkovian (Lower Devonian) (Tarrant, 1991). Palaeoenvironments are generally fluvial, lacustrine or deltaic (Tarrant, 1991).

### Problematica and tesselate heterostracans

A number of mono-generic taxa fall into families of their own, with their affinity to other heterostracans uncertain. Included within the problematic heterostracans are: *Cardipeltis,* an enigmatic heterostracan from the Western-USA, Corvaspididae, *Natlaspis, ?Traquairaspis* (both from the Mackenzie Mountains of Canada) and tessellate heterostracans *Tesseraspis, Lepidaspis, Aporemaspis, Aserotaspis, Kallostrakon* and *Oniscolepis* (the latter two are known only from disarticulated fragments) (Dineley & Loeffler, 1976).

#### Cardipeltis

Three species of *Cardipeltis* have been described from the Pragian & Emsian of the Western-USA (Denison, 1966; Elliott, Reed, & Johnson, 1999). The genus is typified as having a dorso-ventrally flattened carapace with separate dorsal shield and a ventral shield composed of un-fused hexagonal tesserae. Lining the lateral margins are separate plates that are ornamented dorsally and ventrally composing a lateral brim. Unlike other heterostracans, the pair of branchial openings are well inserted into the dorsal shield forming a branchial notch (but not enclosed within it as is seen in some Amphiaspididae). Orbits are not seen in any of the articulated material, nor have any convincing plates been found and the rostral region is seen in one specimen although poorly preserved (Denison, 1966).

Although known from reasonably complete and articulated material, the genus is so unlike other heterostracans that many different hypotheses as to the affinity of *Cardipeltis* have been proposed. These include Cyathaspididae, Pteraspidiformes, Corvaspididae, Psammosteidae and a close affinity to *Kallostrakon* and other tessellate forms (Denison, 1966). *Cardipeltis* is found in the fluvial Water Canyon Formation of Utah and Beartooth Butte Formation of Wyoming dated as Emsian (Denison, 1966; Elliott *et al*., 1999).

#### Corvaspididae

The Corvaspididae contains two genera, namely *Corvaspis* and *Corveolepis. Corvaspis kingi* is the only known species of *Corvaspis* and is regarded as a problematic taxon in part because it is known from a single fragmentary symmetrical plate and a few asymmetrical plates. No obvious dorsal, ventral or branchial plates have been found, however, orbital plates have been assigned to *Corvaspis* material. *Corvaspis* is distinguished by having an ornament pattern uniformly arranged, with smooth topped flat tubercules arranged in tessiform units (Blieck & Karatajūtė-Talimaa, 2001). Also included within Corvaspididae are *Corveolepis arctica* (Loeffler & Dineley, 1976) and *C. elgae* (Blieck & Karatajūtė-Talimaa, 2001). *Corveolepis* has a singular dorsal headshield similar to cyathaspids, with an orbit enclosed within the headshield. The ornament pattern of *Corveolepis* is very similar to *Corvaspis*, hence being united in the same family. However, a fundamental difference between the two taxa is that *Corveolepis* has a dorsal headshield (covering orbital and rostral areas), whereas *Corvaspis* is known from fragmentary separate plates (indicating a multi-plated form).

Tarlo (1960) assigned *Corvaspis* to the psammosteids due to his identification of post-orbital and branchial plates along with the tessellate-pattern seen in their ornament. However, this interpretation cannot be accepted until an articulated specimen is found. There are many differences between *Corvaspis* and the psammosteids, including *Corvaspis’* ornament being different to the psammosteids (which is of stellate tubercules), the ‘tesserae’ identified in *Corvaspis* by Dineley (1953) and Tarlo (1960) is a separate fused lateral plate (Janvier, 1996), and the size and shape of the branchial plate. *Corvaspis* has also been compared to *Cardipeltis*, cyathaspids and as a transitional form between *Kallostrakon*/*Cardipeltis*and the Cyathaspididae (Smith Woodward, 1934; Denison, 1964; Halstead, 1973)*. Corveolepis* and *Corvaspis* have been tentatively united within Corvaspididae based upon similarities in ornament patterns, although the monophyly of this group remains uncertain (Blieck & Karatajūtė-Talimaa, 2001).

*Corvaspis kingi* is known predominantly from the Prìdoli and Lochkovian fluvial deposits of the Welsh Borders, UK (Smith Woodward, 1934; Dineley, 1953, 1965; Tarlo, 1960). *Corveolepis arctica* is described from the Prìdoli of the Canadian Arctic (Loeffler & Dineley, 1976) and *C. elgae* is known from the Lochkovian of Severnaya Zemlya (Blieck & Karatajūtė-Talimaa, 2001).

#### Tesselate heterostracans

Tessellate heterostracans include *Lepidaspis, Aporemaspis* and *Aserotaspis* from Canada and *Oniscolepis, Kallostrakon* and *Tesseraspis* (from modern day Estonia and the UK)(Tarlo, 1964; Halstead Tarlo, 1967; Dineley & Loeffler, 1976; Elliott & Loeffler, 1989; Märss & Karatajūtė-Talimaa, 2009). They are described as tessellate because their dermal armour is composed of smaller subunits or many small platelets. The subunits are generally un-fused but can become fused at the base as is seen in *Lepidaspis* (Dineley & Loeffler, 1976). The affinity of these taxa is problematic as the majority of taxa are only known from fragmentary remains. They are useful for biostratigraphic correlation or as evidence of early vertebrate occurrences, but due to their poor fossil record the anatomy of these taxa remains unknown making them difficult to place both taxonomically and phylogenetically. Despite the tessellate nature of their dermal skeletons, *Aporemaspis, Aserotaspis* and *Lepidaspis* are known from articulated or semi articulated specimens, which preserve key features such as the orbits and pineal regions. However, none of the tessellate heterostracans have their branchial region preserved, making precise affinities more ambiguous. The Ordovician tessellate pteraspidimorph *Astraspis* has multiple branchial openings (Elliott, 1987; Elliott & Loeffler, 1989), and it is not currently clear whether the tessellate heterostracans have a closer affinity to the Ordovician forms or to the Heterostraci. All genera have varying degrees of fusion of the tesserae composing their dermal skeletons. *Aporemaspis* and *Aserotaspis* are both described as having fused regions around their orbits (Dineley & Loeffler, 1976; Elliott & Loeffler, 1989). *Lepidaspis* is probably the best known of the tessellate heterostracans with many articulated specimens. Preservation varies from singular dermal elements (tesserae) to tesserae that have begun to fuse at the boney middle and base layers. *Lepidaspis* has anterior-laterally placed orbits and an anteriorly placed mouth, with a separate dorsal and ventral shield (Dineley & Loeffler, 1976). The holotype of *Tesseraspis* is the most articulated with regions of ‘fused’ tesserae. No orbital, pineal or branchial regions are preserved, but there are two rows of ridged tesserae which have greater relief and coarser tubercules compared the rest of the dorsal shield (similar to *Astraspis*)(Wills, 1935). *Kallostrakon* and *Oniscolepis* are only known from disarticulated fragments of bone and are placed in the Heterostraci mostly due to similarities in histology (Märss & Karatajūtė-Talimaa, 2009).

The affinities of tessellate heterostracans remains unclear. Due to similarities to Ordovician pteraspidimorphs (i.e. sharing with *Astraspis* and *Eriptychius* a dermal shield composed of smaller platelets) the tessellate heterostracans have often been regarded as the plesiomorphic state for the Heterostraci, bridging the gap between the Ordovician forms and the rest of the Heterostraci (‘tessellate-basal’ model) (Tarlo, 1965; Halstead, 1973; Janvier, 1996). However, they have yet to be included in any formal phylogenetic analysis. Tessellate heterostracans are found in a range of localities including the Canadian Arctic and Mackenzie mountain regions, Welsh Borders and Baltic regions. These were contained in the palaeocontinents Laurentia, Baltica and Avalonia and are generally found in fluvial sediments.

#### Natlaspis

*Natlaspis* is a mono-specific genus. Its headshield is composed of dorsal and ventral shields, the dorsal of which encloses the orbital, branchial and pineal organs. Its ornament is of coarse serrated tubercules which are well spaced.Dineley & Loeffler (1976) placed *Natlaspis* in *incertae familiae* although suggested it may belong in a new subfamily with *?Traquairaspis*. They both have similar headshield components and most likely require re-description and elevation to a new family. *Natlaspis* is yielded from the Delorme Formation (Laurentia) dated as Prìdoli (Mackenzie Mountains) and interpreted as hyposaline or a brackish lagoon (Dineley & Loeffler,1976)

#### ?Traquairaspis

Some species from Delorme Formation of the Mackenzie Mountains Canada have uncertain affinity, potentially *Traquairaspis*. They are characterized by having a dorsal shield that encloses their pineal, branchial and orbital openings. The ornament pattern is of highly crenulated discontinuous dentine ridges that sometimes form elevated crests. Specimens belonging to *?Traquairaspis* have been tentatively placed within the Traquairaspididae by Dineley & Loeffler (1976). They have previously not been included in any phylogenetic analysis. All *?Traquairaspis* specimens are from the Laurentian Prìdoli Delorme Formation in the Mackenzie Mountains interpreted as hyposaline or a brackish lagoon (Dineley & Loeffler,1976)

### Non-heterostracan pteraspidimorphs

There are a number of Ordovician pteraspidimorphs known from articulated specimens, including; *Arandaspis, Sacabambaspis, Astraspis and Eriptychius. Sacabambaspis* and *Arandaspis* have large singular dorsal and ventral headshields and a trunk region covered in elongate rod shaped scales (Ritchie & Gilbert-tomlinson, 1977; Gagnier, Blieck, & Rodrigos, 1986; Gagnier, 1995). *Astraspis* and *Eriptychius* are less completely preserved then Arandaspida taxa due to their headshields being composed of non-fused platelets or tesserae (Denison, 1967; Elliott, 1987; Sansom *et al*., 1995; Sansom & Smith, 2005). Unlike the heterostracans, the Ordovician pteraspidimorphs appear to have more than one gill or branchial opening, but histologically are very similar to heterostracans, possessing a three-layered dermal skeleton comprising a superficial layer of dentine tubercles capped with enameloid, a middle vascular layer composed of tissue similar to aspidin and a compact lamellar basal layer (Denison 1967, Sansom, Smith, & Smith, 1997; Sansom, Donoghue, & Albanesi, 2005, Lemierre & Germain 2019). *Sacabambaspis* is an unusual form with its orbits placed at the anterior of its headshield, next to which are another pair of openings (interpreted as external nostril openings) (Gagnier *et al*., 1986). Another unusual feature is a pair of openings placed towards the middle of the dorsal shield interpreted by Gagnier (1995) as a double pineal opening, which is unknown in any other early vertebrates and is potentially homologous with the endolymphatic openings seen in osteostracans.

The Ordovician Pteraspidomorphi have previously been interpreted as separate from the Heterostraci clade due to their lack of a single pair of branchial openings. However, they share many features in common with the Heterostraci including a dermal skeleton composed of aspidin and topped with enameloid and tubular orthodentine (Sansom *et al*., 1997, 2005). They are traditionally placed as sister taxa to the Heterostraci and all are united in the clade Pteraspidimorphi. However, which of the Ordovician forms is the sister group to the Heterostraci remains controversial (i.e. the arandaspids (*Arandaspis* and *Sacabambaspis*) or the tessellate pteraspidimorphs (*Astraspis* and *Eriptychius*)). Some authors hypothesise that the tessellate forms are sister group to the Heterostraci and that *Tesseraspis* and other problematica are plesiomorphic heterostracans (‘tessellate-basal’ theory). The alternative hypothesis is that arandaspids are sister group indicating the plesiomorphic condition for the Heterostraci is a headshield covered in large singular dorsal and ventral plates. It must be noted, however, that the ventral side of *Sacabambaspis* is composed of apparently fused tesserae (Gagnier *et al*., 1986) indicating the large plated forms could have a tessellate affinity.

With respect to their age and location, *Arandaspis* is the oldest articulated vertebrate and is known from the Stairway Sandstone Formation (Lower Ordovician) of Australia (Ritchie & Gilbert-tomlinson, 1977). *Sacabambaspis* is from the Upper Ordovician Anzaldo Formation in Bolivia, both these localities were situated in the super-continent Gondwana (Gagnier *et al*., 1986). *Astraspis* and *Eriptychius* both belong to the Harding Sandstone Formation and Winnipeg Formations of the United States and Canada respectively (Sansom *et al*., 1995; Sansom & Smith, 2005).

## 4. PREVIOUS INTERPRETATIONS OF HETEROSTRACAN RELATIONSHIPS

Only the Pteraspidiformes (Ilyes & Elliott, 1994; Pernègre, 2002; Pernègre & Goujet, 2007; Pernègre & Elliott, 2008; Randle & Sansom, 2016, 2017) and Cyathaspididae heterostracans (Lundgren & Blom, 2013; Randle & Sansom, 2017) have undergone any formal cladistic analysis. However, the inter-relationships of the Heterostraci have been considered by a number of authors, placing them into higher-level taxonomic clades. White (1935) placed them into disparate clades based on similarities in their anatomy. Interestingly he split the Cyathaspididae (*sensu* Denison,1964) into two different groups: one for cyathaspids without epitega (Palaeaspidae including *Poraspis*) and the other with epitega (Cyathaspidae including *Cyathaspis* and *Archegoneaspis*).

Obruchev (1967) was one of the first to consider the intra-relationships of heterostracans (Fig. 4A). He interpreted the Cyathaspididae as paraphyletic with respect to the Amphiaspididae, and the Psammosteidae and Pteraspidiformes as sister groups. His conception of heterostracan relationships did not include the tessellate heterostracans and Traquairaspididae were placed in a polytomy with the remaining heterostracan clades.

Halstead (1973) (Fig. 4B) produced the first tree containing all representatives of the Heterostraci demonstrating his perceived evolutionary relationships for the clade. He interpreted the plesiomorphic headshield condition for the Heterostraci as tessellate, e.g. *Tesseraspis* (being composed of small boney units termed tesserae), derived from the tessellate Ordovician forms e.g. *Astraspis* (tessellate-basal model). Following this scheme, the heterostracans then divided with one lineage leading to *Cardipeltis*, amphiaspids and the cyathaspids and the other psammosteids, traquairaspids and pteraspids with each of these lineages ‘passing through’ partially tessellate forms (*Cardipeltis* and psammosteids respectively).

Novitskaya (1983) (Fig. 4E) considered the relationships of Amphiaspididae, Cyathaspididae and Pteraspidiformes. She believed the Amphiaspididae were derived from the Cyathaspididae as they share a similar ornament pattern and some Amphiaspididae appear cyathaspid-like. For example, the amphiaspid *Aphataspis* has a vestige of the suborbital plate seen in the cyathaspids, and she believes *Boothiaspis* (a cyathaspid) and *Argyriaspis* (an amphiaspid) share features such as their overall dimensions and shape of the dorsal shield and have similarly positioned mouth and orbits. In her figure (text Fig. 76, p163 (Novitskaya, 1983)) of evolutionary relationships she placed *Ctenaspis* as sister group to Amphiaspididae, and reconstructed cyathaspids as paraphyletic and more closely related to the Pteraspidiformes than the Amphiaspididae.

Contra to Novitskaya (1983), Blieck *et al*. (1991) (Fig. 4D) interpreted the Cyathaspididae and Amphiaspididae as one clade (labelled CA clade) and the Pteraspidiformes, Psammosteidae and Anchipteraspididae as another (named APP clade). They defined the CA group as having: longitudinally arranged ornament patterns, flank scales being long and rectangular, absence of looping inter-orbital and the ‘fusion’ of the branchial plate to the dorsal shield and enclosed orbits. Whereas, the APP group are defined by: a continuous inter-orbital canal (pineal of some) around or through the pineal plate, rhombic scales, concentric ornament pattern in the rostral, pineal and dorsal plates and lastly having separate (Blieck *et al*., 1991).

Janvier (1996) interpreted the cyathaspids as the sister group to the Amphiaspididae and ctenaspids. These were united under the Cyathaspidiformes, which is in turn were sister group to the Pteraspidiformes (Psammosteidae + pteraspids). All these were informally labelled the ‘higher heterostracans’ defined by a branchial plate not pierced by the branchial opening (Janvier, 1996). In a polytomy at the base of the heterostracans are *Lepidaspis, Toombsaspis* (a member of the Traquairaspididae) and *?Traquairaspis mackenziensis* (Fig.4C). These taxa are generally regarded as problematic with unknown affinities.

The majority of phylogenetic analyses including heterostracans have been clade specific, or only included one representative taxon from another clade. Randle & Sansom (2017) (Fig. 4F) were the first to include two major clades i.e. all taxonomically described Cyathaspididae and Pteraspidiformes. Their results indicated the Cyathaspididae were paraphyletic with regards to Pteraspidiformes. However, without the inclusion of groups such as the Amphiaspididae and Traquairaspididae in a phylogenetic analysis, the exact relationships of these clades remain ambiguous.

## 5. MATERIALS AND METHODS

### Taxon Sample

#### Ingroup

We took a comprehensive approach to taxon sampling of Heterostraci aiming to include as many genera as possible. Our final sample of 145 ingroup taxa, encompassed the broad taxonomic and morphological diversity seen within the heterostracans. Where possible, the type species of a genus was used, unless another was known from better preserved or articulated material. Character codings are based on the species included as the representative of that genus rather than for a composite of all species within a genus. Two species were included for some genera within the analyses to either test monophyly or because it was deemed inclusion of both held important phylogenetic information. These include; *Ariaspis, Cardipeltis, Pionaspis, Lampraspis, Protopteraspis, Corveolepis* and *Homalaspidella*.

#### Outgroup

Outgroup choice is extremely important for any phylogenetic analysis; this is especially true for the Heterostraci and Pteraspidimorphi. There is no clear consensus as to which of the non-heterostracan pteraspidimorphs are the sister group to the Heterostraci, therefore all i.e. *Astraspis*, *Arandaspis* and *Sacabambaspis* were included in the analysis. The position of the Heterostraci on the gnathostome stem is also ambiguous; therefore representatives of all major plesia on the stem group were included. The phylogenetic matrices of Keating & Donoghue (2016) and Gabbott et al (2016) were used due to wide taxon sampling and characters that were created to discern the phylogenetic relationships between early vertebrate groups. These matrices stem from a number of other phylogenetic works including: Donoghue, Forey, & Aldridge (2000), Gess, Coates, & Rubidge (2006), Sansom *et al*., (2010), Blom (2012) and Gabbott *et al*., (2016).

### Characters

In total, 343 characters were coded. Characters were constructed based on anatomical observations, taxonomic descriptions and sourced from previous phylogenetic analyses, primarily; Ilyes & Elliott (1994); Pernègre (2002); Pernègre & Goujet (2007); Pernègre & Elliott (2008); Lundgren & Blom (2013); Randle & Sansom (2016) and Randle & Sansom (2017) for the Heterostraci. The matrix of Keating & Donoghue (2016) including non-heterostracan taxa was used and expanded.

Of those 343, 25 characters related to quantitative traits that treated discretised characters (Randle & Sansom 2016, 2017). Discretized characters are treated as ordered. Discretized characters were weighted by 1/(*n* −1), when *n*= number of character states (not applicable if only two character states). Two quantitative characters (271 & 289 from the discrete-with-discretised analysis), which have 7 characters states, were amended to only have 6 character states (the maximum for Bayesian software MrBayes). This was achieved by merging character states 0 and 1 of character 271 (as character state 0 contained one taxon and is therefore uninformative) and state 5 and 6 for character 289 (as character 6 also only contained one taxon). This character coding was used for both parsimony and Bayesian analyses.

### Character List

**1.** Pineal macula enclosed in dorsal shield: (0) absent, (1) present. Pineal organ in dorsal shielded forms usually covered by a small dentine unit, sometimes not always visible by a deformation in the surrounding ornament patterns. Character (ch.) 54 of Lundgren & Blom (2013), ch.1 of Randle & Sansom (2017).
**2.** Pineal plate (separate plate): (0) absent, (1) present. Ch.1 of Randle & Sansom (2016), ch.2 of Randle & Sansom (2017).
**3.** Pineal plate enclosed by dorsal plate: (0) absent, (1) present. Pineal plate enclosed by dorsal plate is condition seen in Anchipteraspididae (Elliott, 1984), see Fig. 3. Ch. 3 of Randle & Sansom (2016), ch.3 of Randle & Sansom (2017).
**4.** Pineal plate morphology (for those that have a pineal plate delimited by the median dorsal and rostral plates): (0) triangular, (1) quadrangular, (2) circular/ovate, (3) flat topped ovate (*Doryaspis*), (4) flat bottomed, triangular-hexagonal top (*Phialaspis* & *Toombsaspis*). Ch.4 of Randle & Sansom (2016; 2017).
**5.** Quadrangular pineal plate morphology: (0) rectangular, (1) anterior and posterior edge convex, (2) anterior edge concave, posterior edge convex, (3) posterior edge convex. Ch.5 of Randle & Sansom (2016; 2017).
**6.** Pineal-orbital belt: (0) absent, (1) present. Presence when the orbital and pineal plates are in contact with each other separating the dorsal and rostral plates. Ch.25 of Pernègre (2002), ch.12 Pernègre & Goujet (2007) and ch.6 of Randle & Sansom (2016; 2017).
**7.** Pineal-orbital plate contact: (0) point contact (very small contact, just point to point), (1) side contact (ribbon contact of Novitskaya (1983)), (2) pineal plate forms large part of orbital plate margin (forms that lack a medial orbital process (*Phialaspis* & *Toombsaspis*). Ch.4 of Ilyes & Elliott (1994), ch.26 of Pernègre & Elliott (2008) and ch.7 of Randle & Sansom (2016; 2017).
**8.** Rostral plate (separate plate): (0) absent, (1) present. Rostral plate is a separate plate situated at the anterior end of the headshield. Ch.8 of Randle & Sansom (2016; 2017).
**9.** Pre-oral surface (ornamented medial area on the ventral side of rostral plate): (0) absent, (1) present. Ch.9 of Randle & Sansom (2016; 2017).
**10.** Pre-oral field (unornamented area on the ventral side of rostral plate prior to the ascending lamella): (0) absent, (1) present. Ch.10 of Randle & Sansom (2016; 2017).
**11.** Ornamentation of pre-oral surface: (0) transverse (1) variable orientation of ridges. Ch.11 of Randle & Sansom (2016; 2017).
**12.** Shape of anterior end of rostral plate: (0) rounded, (1) rounded to a point, (2) truncated/ concave, (3) triangular, Adapted from ch.35 of Pernègre & Goujet (2007), ch.30 of Pernègre & Elliott (2008) and ch.12 of Randle & Sansom (2016; 2017).
**13.** Orbital notch in rostral plate: (0) absent, (1) present. Adapted from ch.24 of Pernègre (2002), ch.11 of Pernègre & Goujet (2007) ch.19 of Pernègre & Elliott (2008) and ch.13 of Randle & Sansom (2016; 2017).
**14.** Orbital notch in the rostral plate: (0) rounded, (1) angular. Ch.14 of Randle & Sansom (2016; 2017).
**15.** Rostral plate pineal plate contact: (0) concave contact, (1) no notch, (2) convex notch. Adapted from ch.14 of Pernègre & Goujet (2007), ch.28 of Pernègre & Elliott (2008) and ch.15 of Randle & Sansom (2016; 2017).
**16.** Position of the orbits: (0) lateral, (1) dorsal. Coded as dorsal if the entire orbital opening is seen in dorsal view indicating positioned more on top of the shield. Ch.18 of Randle & Sansom (2016; 2017).
**17.** Orbits delimited by the dorsal shield: (0) absent, (1) present. Ch. 17 of Randle & Sansom (2017).
**18.** Orbits entirely enclosed within dorsal shield: (0) absent, (1) present. Ch. 18 of Randle & Sansom (2017).
**19.** A single pair of branchial plates: (0) absent, (1) present, Some anaspids and heterostracan have a single plate in the branchial region, of is not known f these are homologous. Ch. 3 of Blom 2012, ch.19 of Randle & Sansom (2017).
**20.** Plates covering the branchial region (separate plate): (0) absent, (1) present. Coded absent for taxa whose branchial openings seemingly external open. Ch. 20 of Randle & Sansom (2017).
**21.** Sub-branchial scales: (0) absent, (1) present. Ch.4 of Lundgren & Blom (2013), ch.21 of Randle & Sansom (2017).
**22.** Lateral branchial opening position: (0) lateral side of shield, (1) posterior end of headshield. (Not applicable to taxa whose branchial openings are ventral or dorsal in aspect). Ch.1 of Ilyes & Elliott (1994), ch.5 of Pernègre & Goujet (2007), ch.1 of Pernègre & Elliott (2008) and ch.20 of Randle & Sansom (2016), ch.22 of Randle & Sansom (2017).
**23.** Dorsal plate delimits part of the single pair of branchial openings: (0) absent, (1) present. Contingent on possessing a dorsal plate. Ch. 23 of Randle & Sansom (2017).
**24.** Cornual plate delimits part of the single pair of branchial openings: (0) absent, (1) present. Contingent on possessing a cornual plate. Ch. 24 of Randle & Sansom (2017).
**25.** Dorsal shield delimits part of the single pair of branchial openings: (0) absent, (1) present. Contingent on possessing a dorsal shield. Ch. 25 of Randle & Sansom (2017).
**26.** Single pair of branchial openings bisect the dorsal shield: (0) absent, (1) present. Contingent on dorsal shield delimiting branchial opening. Ch. 26 of Randle & Sansom (2017).
**27.** Single pair of branchial openings bisect the branchial plate: (0) absent, (1) present. Contingent on possessing a branchial plate. Ch. 27 of Randle & Sansom (2017).
**28.** Ventral shield delimits the single pair of branchial openings: (0) absent, (1) present. Contingent on possessing a ventral shield. Ch. 28 of Randle & Sansom (2017).
**29.** Lateral brim delimits part of the single pair of branchial openings: (0) absent, (1) present. Contingent on possessing a lateral brim. Ch. 29 of Randle & Sansom (2017).
**30.** Pair of cornual plates (separate plates): (0) absent, (1) present. Adapted from ch.9 of Pernègre & Goujet (2007), ch.12 of Pernègre & Elliott (2008) and ch.21 of Randle & Sansom (2016) and ch.30 of Randle & Sansom (2017).
**31.** General cornual plate morphology: (0) lateral (external) and posterior sides convex, (1) lateral (external) side concave and posterior side convex (*Doryaspis*), (2) lateral (external) side convex and posterior side concave, (3) all sides rounded/convex so appears triangular (see Drepanaspis). Ch.22 of Randle & Sansom (2016) and ch.31 of Randle & Sansom (2017).
**32.** Lateral projection of cornual plates: (0) lateral projection or cornual plate less than branchial plate, (1) lateral projection the same as branchial plates, (2) lateral projection of cornual just up to greater than double that of the branchial plates (3) lateral projection vastly greater than branchial plate (4) cornual plates placed on dorsal shield. Ch.23 of Randle & Sansom (2016) and ch.32 of Randle & Sansom (2017). [Ordered]
**33.** Posterior extension of cornual plates: (0) less than posterior margin of dorsal plate, (1) equal to posterior margin of dorsal plate, (2) greater than posterior margin of dorsal plate. Ch.24 of Randle & Sansom (2016) and ch.33 of Randle & Sansom (2017). [Ordered]
**34.** Ornamentation of cornual plates: (0) scale like ornamentation, (1) long ridges parallel to the lateral (external) edge (3) concentric ornament (Psammosteids – need to figure out what to do about these characters). Ch. 25 of Randle & Sansom (2016) and ch.34 of Randle & Sansom (2017).
**35.** Dorsal plate (medial, separate): (0) absent, (1) present. Ch. 35 of Randle & Sansom (2017).
**36.** Dorsal plate surrounded by ‘fields of tesserae’ or platelets: (0) absent, (1) present. Ch. 36 of Randle & Sansom (2017).
**37.** Posterior margin of dorsal plate: (0) sinuous medial peak, (1) medial peak, (2) straight, (3) sinuous, (4) rounded, (5) notched as seen in dorsal view. Adapted ch.27 of Pernègre & Goujet (2007), ch.35 of Pernègre & Elliott (2008), ch.29 of Randle & Sansom (2016) and ch.37 of Randle & Sansom (2017).
**38.** Branchial notch in dorsal plate: (0) absent, (1) present. Notch in the dorsal shield for branchial opening. Ch.30 of Randle & Sansom (2016) and ch.38 of Randle & Sansom (2017).
**39.** Embayment in dorsal plate to accommodate the cornual plates: (0) absent, (1) present (dorsal flexure of Voichyshyn 2011). Ch.31 of Randle & Sansom (2016) and ch.39 of Randle & Sansom (2017).
**40.** Notch at posterior end of the dorsal plate (to house the dorsal spine, can be enclosed): (0) absent, (1) present. Ch.33 of Randle & Sansom (2016) and ch.40 of Randle & Sansom (2017).
**41.** Dorsal shield: (0) absent, (1) present. The dorsal shield is generally the only shield present in the dorsal part of the headshield. It encompasses the rostral and orbital areas and is a single element (different from that in Pteraspidiformes, in which the dorsal headshield is composed of many different plates. Ch. 41 of Randle & Sansom (2017).
**42.** Maxillary brim: (0) absent, (1) present. Ventral fold at anterior end of dorsal shield, different to the pre-oral field and surface of Pteraspidiformes which is on the ventral side of the rostral plate. Ch. 42 of Randle & Sansom (2017).
**43.** Ornamented maxillary brim: (0) absent, (1) present. Ch. 43 of Randle & Sansom (2017).
**44.** Medial rostral process: (0) absent, (1) present. Distinct medial fold seen at anterior end or dorsal shield (rostral area). Ch.10 of Lundgren & Blom (2013) and ch.44 of Randle & Sansom (2017).
**45.** Pre-orbital process: (0) absent, (1) present. Downward folding of dorsal shield (cyathaspids) forming the anterio-dorsal margin of the orbital opening. Contingent on having the orbital opening surrounded by the dorsal shield dorsally and either the ventral shield or sub-orbital plate ventrally. Ch. 45 of Randle & Sansom (2017).
**46.** Pre-orbital suture: (0) absent, (1) present. Break in ornament preceding or on the pre-orbital process in taxa with a dorsal shield, seen in *Alainaspis* and *Boothiaspis* (Elliott & Dineley, 1985). Ch. 46 of Randle & Sansom (2017).
**47.** Supra-orbital arch/crest on dorsal shield: (0) absent, (1) present. Seen in *Boothiaspis* and *Alainaspis* (Broad, 1973; Elliott & Dineley, 1985, 1991). Ch. 47 of Randle & Sansom (2017).
**48.** Post-branchial lobes: (0) absent, (1) present. Downward folding of the dorsal shield posterior to the branchial opening. Adapted from ch.20 of Lundgren & Blom (2013) and ch.48 of Randle & Sansom (2017).
**49.** Lateral brim: (0) absent, (1) present. Ch.14 of Lundgren & Blom (2013) and ch. 49 of Randle & Sansom (2017). Lateral brims are a lateral extension of the dorsal shield, generally with a different relief to that of the medial area on the dorsal shield, different to epitega which are divisions based upon ornament.
**50.** Serrated lateral brim: (0) absent, (1) present. Ch.15 of Lundgren & Blom (2013) and ch.50 of Randle & Sansom (2017).
**51.** Pointed posterior end of lateral brim: (0) absent, (1) present. Character 22 Lundgren & Blom (2013) and ch.51 of Randle & Sansom (2017).
**52.** Lateral lamina on the dorsal shield (downwardly projecting lamina): (0) absent, (1) present. Ch. 17&18 of Lundgren & Blom (2013) and ch.52 of Randle & Sansom (2017).
**53.** Dorsal spine: (0) absent, (1) present. Adapted from ch.2 of Pernègre (2002), ch.10 of Pernègre & Elliott (2008) and ch.34 of Randle & Sansom (2016) and ch.53 of Randle & Sansom (2017).
**54.** Separate dorsal spine: (0) absent, (1) present. Dorsal spine is its own separate plate not fused to the dorsal shield as in some Cyathaspididae. Ch.35 of Randle & Sansom (2016) and ch.54 of Randle & Sansom (2017).
**55.** Dorsal spine base enclosed by dorsal plate: (0) absent, (1) present. Ch. 36 of Randle & Sansom (2016) and ch.55 of Randle & Sansom (2017).
**56.** Ornamentation of dorsal spine: (0) scale like, (1) longitudinal ridges. Ch. 37 of Randle & Sansom (2016) and ch.56 of Randle & Sansom (2017).
**57.** Posterior medial crest (keel of some authors)(posterio-median crest): (0) absent, (1) present. Different from the dorsal spine as it appears to be a continuation of the dorsal shield. Ch.17 of Lundgren & Blom (2013) and ch.57 of Randle & Sansom (2017).
**58.** Dorsal median medial crest (keel of some authors): (0) absent, (1) present. As seen in *Liliaspis* not to be confused with the crest at the posterior end, this one is more central see character 57. Ch. 58 of Randle & Sansom (2017).
**59.** Medial dorsal peak/process: (0) absent, (1) present. This is a posteriorly orientated extension/peak of the dorsal plate/shield (*Doryaspis*, *Xylaspis*, and some cyathaspids). Ch.32 of Randle & Sansom (2016) and ch.59 of Randle & Sansom (2017).
**60.** Orientation of the mouth: (0) ventral position, (1) dorsal position (2) anterior. Ch.1 of Pernègre (2002) and ch.38 of Randle & Sansom (2016) and ch.60 of Randle & Sansom (2017).
**61.** Number of pairs of lateral post-oral (orogonal) plates: (0) 1 pair, (1) 2 pairs. Ch.8 of Pernègre (2002), ch.53 of Pernègre & Elliott (2008) and ch.39 of Randle & Sansom (2016) and ch.61 of Randle & Sansom (2017).
**62.** Accessory plates along dorsal plate margin: (0) absent, (1) present. Ch.6 of Ilyes & Elliott (1994), ch.27 of Pernègre (2002), ch.5 of Pernègre & Elliott (2008) and ch.40 of Randle & Sansom (2016) and ch.62 of Randle & Sansom (2017).
**63.** Body scales (relative to trunk region): (0) deep and rectangular, (1) small and diamond shaped. Ch.63 of Randle & Sansom (2017).
**64.** Finer scales in between digitations in caudal fin: (0) absent, (1) present. Ch.5 of Pernègre (2002) and ch.64 of Randle & Sansom (2017). Absent of these features as seen in *Nahanniaspis* which is described as having a forked tail.
**65.** Caudal fin morphology: (0) ventral lobe more pronounced than dorsal, (1) dorsal and ventral lobe equally developed, (2) dorsal lobe more pronounced than the ventral, (3) no distinct lobes. Ch. 65 of Randle & Sansom (2017).
**66.** Internal organ impression on the visceral side of the dorsal plate/shield (only applicable to taxa with a singular dorsal head plate or shield): (0) absent, (1) present. Ch.38 of Pernègre & Goujet (2007), ch.36 of Pernègre & Elliott (2008) ch.41 of Randle & Sansom (2016) and ch.66 of Randle & Sansom (2017).
**67.** Continuous tubercle ridges (ornamentation) pattern on dorsal shield/plate: (0) absent, (1) present. Condition for Pteraspidiformes and some Cyathaspididae, but not *Drepanaspis*. Some psammosteid taxa have a mixture of stellate and ridges (where stellate tubercules appear to have merged), it is therefore not a multistate character. Adapted from ch.43 of Randle & Sansom (2016) and ch.67 of Randle & Sansom (2017).
**68.** Arrangement/deposition of dentine tubercle on dorsal plate/shield as seen at the central anterior end of dorsal shield: (0) lateral addition of dentine ridges, (1) concentric circles from a primordium, (2) radiating from a primordium. Relating to pattern of dentine ridges and associated growth of dentine tubercules. Pteraspidiformes and some cyathaspids have a pattern of seemingly concentric ridges, it is unknown if the Pteraspidiformes pattern is homologous to that of the cyathaspids. In the cyathaspids, coding is for those with of without epitega. For those with epitega it relates to the pattern on the central epitega and those with a post-rostral field relates to the general pattern perhaps behind this. Whorled irregular pattern seen in *Nahanniaspis* and *Torpedaspis*, their pattern is more irregular than other groups and cant be placed clearly into either of the other two groups. Adapted from ch.44 of Randle & Sansom (2016) and ch.68 of Randle & Sansom (2017).
**69.** Position of primordium for concentric growth on dorsal plate/shield: (0) primordium in the anterior half of the dorsal plate/shield and midline (1) primordium in the center of dorsal plate, (2) primordium in the posterior half of the dorsal plate. Contingent on character 65, state (1). Adapted from ch.11 of Pernègre (2002), ch.7 of Pernègre & Goujet (2007), ch.8 of Pernègre & Elliott (2008) ch.45 of Randle & Sansom (2016) and ch.69 of Randle & Sansom (2017). [Ordered]
**70.** Tubercle edge ornamentation (tubercules from dorsal shield/plate can be taxa with dermal skeleton topped with dentine and enameloid of take whose ornament is of the reticulate layer): (0) smooth ridges, (1) crenulate ridges, (2) serrated, (3) scalloped. Adapted from ch.21 of Pernègre & Goujet (2007), ch.9 of Pernègre & Elliott (2008) ch.46 of Randle & Sansom (2016) and ch.70 of Randle & Sansom (2017).
**71.** Tops of dentine ridges (tubercules from dorsal shield/plate): (0) smooth/rounded, (1) crested, (2) flat. Ch.47 of Randle & Sansom (2016) and ch.71 of Randle & Sansom (2017).
**72.** Ornament configuration within the banding (for taxa with ridges): (0) uniform (continuous ridges), (1) undulating ridges (tuberculated ridges), (2) rows of tubercles. Adapted from ch.2 of Ilyes & Elliott (1994) ch.48 of Randle & Sansom (2016) and ch.72 of Randle & Sansom (2017).
**73.** Anterior rostral ridge pattern for taxa with a dorsal shield: (0) transverse, (1) radiating. Ch. 51 & 52 of Lundgren & Blom (2013) and ch.73 of Randle & Sansom (2017). Independent of whether the taxa have rostral epitega.
**74.** Pineal triangle/post-rostral field: (0) absent, (1) present. Distinct triangle area (change in ornament flow or type) at the anterior end of the shield radiating from the pineal forwards, usually demarked by a change in ornamentation associated with the SOC ca1al (Denison, 1964). Adapted from ch.48 of Lundgren & Blom (2013) and ch.74 of Randle & Sansom (2017).
**75.** Dorsal ornament contains superficial scale like units (small concentric esq. units): (0) absent, (1) present. Ornament of smaller scale like ridges. Adapted from ch.39 of Lundgren & Blom (2013) and ch.75 of Randle & Sansom (2017).
**76.** Coverage of scale like units on dorsal shield: (0) majority shield, (1) posterior end only, (2) arranged in rows along flanks of shield. Adapted from ch.40 of Lundgren & Blom (2013) and ch.76 of Randle & Sansom (2017).
**77.** Differing height/width of ridges on dorsal shield/plate: (0) absent, (1) present. This could be in scale like units e.g. *Tolypelepis* or taxa such as *Cyathaspis* where there is a seemingly central wider, higher dentine unit surrounded by smaller ones, ch.77 of Randle & Sansom (2017).
**78.** Epitega: (0) absent, (1) present. Adapted from ch.24 Lundgren & Blom (2013) and ch.78 of Randle & Sansom (2017). Dorsal shield is divided into sub-units based upon differences in the ornament pattern. Divided into the central epitegum, rostral and paired lateral epitega. This subdivision is seen in a number of heterostracan taxa including *?Traquairaspis*, some cyathaspids and some amphiaspids.
**79.** Orbital epitegum: (0) absence, (1) presence. Ch.79 of Randle & Sansom (2017).
**80.** Rostral and lateral epitega relief compared to the central epitega: (0) flattened relief compared to the central epitega, (1) no difference in relief (continuation form the central epitega), (2) enlarged relief (“too big shoe” of Lundgren & Blom (2013) etc) (3) rostral has no difference in relief but lateral are flattened. Adapted from ch.12&13 of Lundgren & Blom (2013) and ch.80 of Randle & Sansom (2017).
**81.** Inter-epitegal band of tubercles (dorsal shield): (0) absent, (1) present. Adapted from Ch.38 of Lundgren & Blom (2013) and ch.81 of Randle & Sansom (2017). Band or smaller tubercles lining the different epitega areas, contingent on having epitega.
**82.** Conspicuous pores: (0) absent, (1) present. Coded as present if pores are visible as breaks in the surface ornament. Adapted from ch.57 of Lundgren & Blom (2013) and ch.82 of Randle & Sansom (2017).
**83.** Pores specifically surrounded by ornament: (0) absent, (1) present. In some specimens the ornament (ridges) specifically surround/outline the pore rather than just appearing as a break. Ch. 83 of Randle & Sansom (2017).
**84.** Highly branched canal pattern: (0) absent, (1) present. Adapted from ch.61 of Lundgren & Blom (2013) and ch.84 of Randle & Sansom (2017). Described by Blieck & Heintz (1983) as a network or branched canals.
**85.** Anterior end of supra-orbital canal (SOC): (0) convergent, (1) parallel, (2) divergent. Ch.49 of Randle & Sansom (2016) and ch.85 of Randle & Sansom (2017).
**86.** Middle/Posterior end of SOC: (0) convergent, (1) parallel, (2) divergent. Adapted from ch.33/34 of Pernègre & Goujet (2007) and ch.49/50 of Pernègre & Elliott (2008). Ch.50 of Randle & Sansom (2016) and ch.86 of Randle & Sansom (2017).
**87.** Position of SOC on rostral plate (only applicable to taxa that have a rostral plate): (0) medial, (1) lateral. Ch.52 of Randle & Sansom (2016) and ch.87 of Randle & Sansom (2017).
**88.** Posterior end of inter-orbital (pineal canal of others) canal when ends do not meet posteriorly to form the pineal canal (or posterior end of SOC for those without a CM.SO canal): (0) ends before meeting another canal, (1) joins/meets another canal (MDC). Ch. 88 of Randle & Sansom (2017).
**89.** Presence of the CM.SO: (0) absent, (1) present. Canal joining the IFC canal to the SOC or pineal canal (Voichyshyn 2011). Ch. 89 of Randle & Sansom (2017).
**90.** Point where the SOC meets the CM.SO and inter-orbital canals: (0) between orbital and pineal plates, (1) on the pineal plate, (2) on orbital plate, (3) on the rostral plate. Adapted from ch.20 of Pernègre & Goujet (2007), ch.48 of Pernègre & Elliott (2008), ch.51 of Randle & Sansom (2016) and ch.90 of Randle & Sansom (2017).
**91.** Anterior extension of MDC when meeting the pineal canal: (0) pineal plate, (1) orbital plate, (2) meets pineal canal on the dorsal plate, (3) between orbital and pineal plates. Adapted from ch.19 of Pernègre & Goujet (2007), ch.45 of Pernègre & Elliott (2008), ch.53 of Randle & Sansom (2016) and ch.91 of Randle & Sansom (2017).
**92.** Inter-orbital posterior ends join to make a continuous/single canal: (0) absent, (1) present. The pineal canal is a canal that loops through or round the pineal organ. It perhaps is derived from the uniting of the posterior ends of the SOC canal. This canal is not seen in the Cyathaspididae. Adapted from ch.13 of Pernègre (2002), ch.51 of Pernègre & Elliott (2008), ch.56 of Randle & Sansom (2016) and ch.92 of Randle & Sansom (2017).
**93.** Inter-orbital: (0) loops around pineal plate/macula, (1) loops through pineal plate. Inapplicable for taxa in which the posterior ends of the inter-orbital/pineal canal are not joined/converged. This condition was used by Blieck (1984) to distinguish between the Protaspididae and Pteraspididae, but it is now seen in other groups. Adapted ch.5 of Ilyes & Elliott (1994), ch.13 of Pernègre (2002), ch.2 of Pernègre & Goujet (2007), ch.52 of Pernègre & Elliott (2008), ch.57 of Randle & Sansom (2016) and ch.93 of Randle & Sansom (2017).
**94.** Anterior end of MDC when meeting another canal: (0) meets CM.SO, (1) meets inter-orbital canal (2) junction between CM.SO and inter-orbital canal (apparent extension of the SOC canal). Only applicable for taxa where the canal meets another canal. Ch. 94 of Randle & Sansom (2017).
**95.** Anterior MDC begins on the dorsal disc without connection to any other canals: (0) absence, (1) presence. Ch.55 of Randle & Sansom (2016) and ch.95 of Randle & Sansom (2017).
**96.** LDC anterior prolongation: (0) not connected to another canal, (1) connected to another canal. The LDC runs down the lateral sides of the dorsal shield and plates and generally the anterior end bends round tor joins another canal at the posterior end of the orbital opening, this then becomes the IFC. Ch.96 of Randle & Sansom (2017).
**97.** Number of transverse commissures (TC) on the dorsal plate: (0) 3 (1) 4, (2) 1-2. Adapted ch.3 of Pernègre (2002), ch.58 of Randle & Sansom (2016) and ch.97 of Randle & Sansom (2017).
**98.** TC pattern: (0) parallel, (1) radial. Radial condition seen in Pteraspidiformes, parallel in Cyathaspididae and other clades and other condition seen in the Anchipteraspididae. Adapted from ch.39 of Pernègre & Elliott (2008), ch.59 of Randle & Sansom (2016) and ch.98 of Randle & Sansom (2017).
**99.** Pattern of first pair or dorsal transverse commissures in radial forms: (0) straight, (1) concave, (2) convex. Pernègre & Goujet (2007) ch.29, ch.60 of Randle & Sansom (2016) and ch.99 of Randle & Sansom (2017). Only applicable to taxa with 3 pairs of transverse commissures.
**100.** Pattern of second pair or dorsal transverse commissures in radial forms: (0) straight, (1) concave, (2) convex. Ch.30 Pernègre & Goujet (2007), ch.61 of Randle & Sansom (2016) and ch.100 of Randle & Sansom (2017). Only applicable to taxa with 3 pairs of transverse commissures.
**101.** Pattern of third pair or dorsal transverse commissures in radial forms: (0) straight, (1) concave, (2) convex. Pernègre & Goujet (2007) ch.31, ch.62 of Randle & Sansom (2016) and ch.101 of Randle & Sansom (2017). Only applicable to taxa with 3 pairs of transverse commissures.
**102.** Two pairs of continuous median transverse commissures in radial forms: (0) absence, (1) presence.Ch.63 of Randle & Sansom (2016) and ch.102 of Randle & Sansom (2017).
**103.** First pair of median commissures in radial forms: (0) anterior to transverse commissure, (1) continuous from transverse commissures, (2) posterior to transverse commissure. Adapted from ch.64 of Randle & Sansom (2016) and ch.103 of Randle & Sansom (2017). [Ordered]
**104.** Middle transverse commissure contact with LDC in relation to contact with MDC in radial canal forms: (0) anterior, (1) parallel, (2) posterior. When radial the anterior and posterior transverse commissures are always anterior and posterior (respectively) in contact. Ch.65 of Randle & Sansom (2016) and ch.104 of Randle & Sansom (2017). [Ordered]
**105.** Anterior transverse commissure contact with LDC in radial canal forms: (0) anterior third or LDC, (1) median third of LDC. Character 22 taken from Pernègre & Goujet (2007) and ch.66 of Randle & Sansom (2016).
**106.** Ventral plate (separate): (0) absent, (1) present.
**107.** Ventral plate with a posterior notch: (0) absent, (1) present
**108.** Ventral plate posterior notch in-filled (by superficial tesserae): (0) absent, (1) present.
**109.** Superficial tesserae on ventral plate: (0) absent, (1) present.
**110.** Superficial tesserae on dorsal plate: (0) absent, (1) present.
**111.** Superficial tesserae on rostral plate: (0) absent, (1) present.
**112.** Post-orbital plates: (0) absent, (1) present.
**113.** Superficial tesserae on the post-orbital plate: (0) absent, (1) present.
**114.** Rostral plate pineal plate contact: (0) absent, (1) present.
**115.** Ventral plate with a smooth (ornament) central area: (0) absent, (1) present. This is a defining character of the Traquairaspididae, not to be confused with the smooth area on the ventral shield of psammosteids, which most likely formed due to abrasion.
**116.** Dorsal ornament contains a row of concentric tubercule units with a larger central tubercule surrounded by smaller tubercules with lower relief (as seen in traquairaspids): (0) absent, (1) present.
**117.** Ornament on lateral and anterior edges of dorsal plate different to that of the central region: (0) absent, (1) present.
**118.** Ventral ornament contains rows of concentric tubercules (same as ch.116): (0) absent, (1) present.
**119.** Headshield composed of one singular plate encompassing the dorsal, ventral and lateral areas of headshield: (0) absent, (1) present. This is a singular plate that encapsulates the dorsal and ventral side of the head region of the fish, condition seen in amphiaspids (Novitskaya, 2004).
**120.** External orbital opening: (0) absent, (1) present. *Lecaniaspis* and *Eglonaspis* lacked these and were apparently blind (Novitskaia, 2010).
**121.** Tubular mouth: (0) absent, (1) present, Novitskaya (2004), some specimens such as *Eglonaspis, Lechaniaspis* and *Empedaspis* have an elongated mouth tube which is the anterior extension of the head shield. Elongated post-orbital area (Novitskaya, 1983).
**122.** Sensory grooves: (0) absent, (1) present. Sensory grooves are seen as unornamented lines through the superficial ornament that generally cut across the ornament pattern (amphiaspids and psammosteids).
**123.** Sensory grooves lined by dentine ridges: (0) absent, (1) present. In some taxa specimens the sensory grooves are bordered by dentine ridges following the length of the sensory groove, generally cuts across ornament patterns.
**124.** Post-orbital opening: (0) absent, (1) present. The pre-spiracle opening of Javier (1996), generally positioned posterior-laterally to the orbits of some amphiaspids (Novitskaya, 1983).
**125.** Branchial opening position (0) lateral position, (1) dorsal (medial position or centralized. In some amphiaspid forms the paired branchial openings appear to have shifted to a posterior mid-line position e.g. *Edaphaspis*), (2) ventral.
**126.** Ventral medial crest: (0) absent, (1) present. Seen in some amphiaspids and a species of *Arctictenaspis*.
**127.** Serration of lateral brim commences: (0) anterior half of brim, (1) posterior half of brim.
**128.** Anterior of lateral brim commences at anterior end of shield: (0) absent, (1) present.
**129.** Posterior of lateral brim ends before the lateral posterior margin of main shield: (0) absent, (1) present.
**130.** Bone: (0) absent, (1) present. Ch.4 of Keating & Donoghue (2016).
**131.** Calcified dermal skeleton: (0) absent, (1) present. Ch.9 of Keating & Donoghue (2016).
**132.** Dermal skeleton contains/composed of acellular bone: (0) absent, (1) present. Found in many of the ostracoderm groups and some gnathostomes.
**133.** Dermal skeleton contains/composed of Aspidin: (0) absent, (1) present. Ch.14 of Keating & Donoghue (2016).
**134.** Dermal skeleton contains/composed of cellular bone (osteocyte lacunae): (0) absent, (1) present.Some groups such as the Osteostraci contains a mix of acellular and cellular bone. Ch.13 of Keating & Donoghue (2016).
**135.** Perichondral bone: (0) absent, (1) present. Ch.5 of Keating & Donoghue (2016).
**136.** Dentine: (0) absent, (1) present. Ch. 2 of Keating & Donoghue (2016).
**137.** Dermal bone topped with superficial dentine: (0) absent, (1) present. This is not seen in *Obruchevia* (Psammosteidae) and Ctenaspis (Cyathaspididae). Ch.10 of Keating & Donoghue (2016).
**138.** Dentine type in dermal skeleton: (0) mesodentine, (1) orthodentine. Ch.11 of Keating & Donoghue (2016).
**139.** Secondary plueromic dentine: (0) absent, (1) present. Some psammosteids lack enameloid and instead have hyper-mineralised dentine in the form of plueromic dentine (Keating *et al*., 2015; Glinskiy & Mark-kurik, 2016).
**140.** Enamel(oid): (0) absent, (1) present. Ch.3 of Keating & Donoghue (2016).
**141.** Dermal bone topped with enameloid: (0) absent, (1) present This is not seen in *Obruchevia* (Psammosteidae) and Ctenaspis (Cyathaspididae), nor some of the Ordovician Pteraspidimorphi. Ch.12 of Keating & Donoghue (2016).
**142.** Dermal bone contains a dense laminated layer (L3 of Keating et al. 2015): (0) absent, (1) present. Seen at the base of heterostracan dermal bone, a similar type is seen in thelodonts but may not be composed of aspidin. Ch.19 of Keating & Donoghue (2016).
**143.** Vascular spaces in middle of dermal skeleton (L2 of Keating et al. 2015): (0) absent, (1) present. Ch.17 of Keating & Donoghue (2016).
**144.** General morphology of vascular spaces within aspidin bone: (0) single layer of hexagonal/polygonal spaces (Honeycomb), (1) anatomising rounded cell spaces (cancellous). General condition seen in Pteraspidiformes and Cyathaspididae is single layer, various groups have different morphologies.
**145.** Dermal bone contains a reticulate layer (L1 of Keating *et al*. (2015),compact acellular bone below the tubercle layer): (0) absent, (1) present.
**146.** Orbital plate medial process shape (end): (0) pointed (1) present.
**147.** Round/ovate tubercules interspersed between ridges (not same as those associated with sensory pores): (0) absent, (1) present. Many different amphiaspid taxa have circular/ovate tubercules interspersed between their ridge ornament.
**148.** Rounded/ovate tubercules have greater relief than the surrounding ornament: (0) absent, (1) present.
**149.** Coverage of rounded/ovate tubercules: (0) concentrated at the front of the shield, (1) coverage all over shield, (2) coverage predominantly to the posterior of the shield
**150.** Oral platelets: (0) absent, (1) present. Small peg like platelets anterior to the orogonal plates seen in some Pteraspidiformes taxa.
**151.** Orogonal plates: (0) absent, (1) present.
**152.** Medial oral plate: (0) absent, (1) present.
**153.** Ornament as stellate tubercules: (0) absent, (1) present. Stellate tubercules are generally rounded tubercule units as seen in psammosteids
**154.** Dermal skeleton visible between tubercules: (0) absent, (1) present.
**155.** Tubercule ridges (tubercules from dorsal shield/plate): (0) long and continuous, (1) short and discontinuous.
**156.** Teardrop shaped tubercules: (0) absent, (1) present.
**157.** Sensory canals discontinuous: (0) absent, (1) present. In some Cyathaspididae and Amphiaspididae the sensory canals are not continuous but rather made of shorter unconnected lengths (still appear in the pattern of other groups). Adapted from ch.60 of Lundgren & Blom (2013).
**158.** Headshield covered in micromeric dermal armour with no distinction between headshield and trunk regions: (0) absent, (1) present. Adapted ch.113 of Keating & Donoghue 2016.
**159.** Headshield composed of dermal plates differentiated into dorsal and ventral plates (either singular or a composite of plates): (0) absent, (1) present.
**160.** Dorsal headshield partially or as whole composed of tesserae: (0) absent, (1) present
**161.** Ventral shield partially or as whole composed of tesserae: (0) absent, (1) present
**162.** Evidence of fusion of tesserae (in *Lepidaspis* and Osteostrcans this is in the basal layer of the dermal skeleton): (0) absent, (1) present.
**163.** Single pair of branchial plates bisected by multiple branchial openings: (0) absent, (1) present. Condition seen in anaspids with a single pair of branchial plates
**164.** Orbits delimited by orbital plate/orbital-cornual plate: (0) absent, (1) present
**165.** Orbital-cornual plate: (0) absent, (1) present
**166.** Boney headshield covers dorsal and ventral aspects of head with a hole in ventral aspect: (0) absent, (1) present.
**167.** Supra orbital canal: (0) absent, (1) present. Supraorbital canal generally begins anterior to the orbits and passes between them.
**168.** Medial dorsal canal: (0) absent, (1) present.
**169.** Lateral dorsal canal: (0) absent, (1) present.
**170.** Infra-orbital canal: (0) absent, (1) present.
**171.** Olfactory peduncles: (0) absent, (1) present. Ch.2 of Keating & Donoghue (2016).
**172.** Adenohypophysis: (0) absent, (1) present. Ch.4 of Keating & Donoghue (2016).
**173.** Optic tectum: (0) absent, (1) present. Ch.6 of Keating & Donoghue (2016).
**174.** Cerebellar primordia: (0) absent, (1) present. Ch.7 of Keating & Donoghue (2016).
**175.** Flattened spinal chord: (0) absent, (1) present. Ch. 9 of Keating & Donoghue (2016).
**176.** Ventral and dorsal spinal nerve roots united: (0) absent, (1) present. Ch.10 of Keating & Donoghue (2016).
**177.** Mauthner fibers in central nervous system: (0) absent, (1) present. Ch.11 of Keating & Donoghue (2016).
**178.** Retina: (0) absent, (1) present. Ch.12 of Keating & Donoghue (2016).
**179.** Olfactory organ with external opening: (0) absent, (1) present. Ch.13 of Keating & Donoghue (2016).
**180.** Nasohypophyseal opening serving respiration (nasohypophyseal duct): (0) absent, (1) present. Ch. 14 of Keating & Donoghue (2016).
**181.** Otic capsule anterior to branchial series: (0) absent, (1) present. Ch.19 of Keating & Donoghue (2016).
**182.** Semicircular canals in labyrinth: (0) absent, (1) present. Ch.20 of Keating & Donoghue (2016).
**183.** Vertical semicircular canals forming loops that are separate from the roof of the utriculus: (0) absent, (1) present. Ch.21 of Keating & Donoghue (2016).
**184.** Externally open endolymphatic ducts: (0) absent, (1) present. Ch.22 of Keating & Donoghue (2016).
**185.** Electroreceptive cells: (0) absent, (1) present. Ch.23 of Keating & Donoghue (2016).
**186.** Sensory lines: (0) absent, (1) present. Ch.24 of Keating & Donoghue (2016).
**187.** Sensory lines on: (0) head only, (1) on head plus body. Ch.25 of Keating & Donoghue (2016).
**188.** Pouch-shaped gills: (0) absent, (1) present. Ch.27 of Keating & Donoghue (2016).
**189.** Single confluent branchial opening: (0) absent, (1) present. Ch.28 of Keating & Donoghue (2016).
**190.** Elongate branchial series: (0) more than 10 gill pouches/slits, (1) fewer than 10. Ch. 29 of Keating & Donoghue (2016).
**191.** Gill openings lateral and arranged in slanting row: (0) absent, (1) present. Ch.30 of Keating & Donoghue (2016).
**192.** Opercular flaps associated with gill openings: (0) absent, (1) present. Ch.32 of Keating & Donoghue (2016).
**193.** Endodermal gill lamellae: (0) absent, (1) present. Ch.33 of Keating & Donoghue (2016).
**194.** Gill lamellae with filaments: (0) absent, (1) present. Ch.34 of Keating & Donoghue (2016).
**195.** Velum: (0) absent, (1) present. Ch. 37 of Keating & Donoghue (2016).
**196.** Multi-chamber heart: (0) absent, (1) present. Ch.38 of Keating & Donoghue (2016).
**197.** Closed pericardium: (0) absent, (1) present. Ch. 39 of Keating & Donoghue (2016).
**198.** Open blood system: (0) absent, (1) present. Ch.40 of Keating & Donoghue (2016).
**199.** Paired dorsal aortae: (0) absent, (1) present. Ch.41 of Keating & Donoghue (2016).
**200.** Large lateral head vein: (0) absent, (1) present. Ch. 42 of Keating & Donoghue (2016).
**201.** Lymphocytes: (0) absent, (1) present. Ch.43 of Keating & Donoghue (2016).
**202.** Subaponeurotic vascular plexus: (0) absent, (1) present. Ch.44 of Keating & Donoghue (2016).
**203.** Dorsal fin: separate dorsal fin: (0) absent, (1) present. Ch.45 of Keating & Donoghue (2016).
**204.** Anal fin separate: (0) absent, (1) present. Ch.47 of Keating & Donoghue (2016).
**205.** Fin ray supports: (0) absent, (1) present. Ch.48 of Keating & Donoghue (2016).
**206.** Paired antero-posterior skin folds: (0) absent, (1) present. Ch.49 of Keating & Donoghue (2016).
**207.** Constricted pectoral fins with endoskeletal elements: (0) absent, (1) present. Ch.50 of Keating & Donoghue (2016).
**208.** Chordal disposition relative to tail development: (0) isochordal, (1) hypochordal, (2) hyperchordal. Ch.53 of Keating & Donoghue (2016).
**209.** Pre-anal median fold: (0) absent, (1) present. Ch.54 of Keating & Donoghue (2016).
**210.** Ability to synthesize creatine phosphatase: (0) absent, (1) present. Ch.55 of Keating & Donoghue (2016).
**211.** Visceral arches fused to the neurocranium: (0) absent, (1) present. Ch.56 of Keating & Donoghue (2016).
**212.** Keratinous teeth: (0) absent, (1) present. Ch.57 of Keating & Donoghue (2016).
**213.** Trematic rings: (0) absent, (1) present. Ch.60 of Keating & Donoghue (2016).
**214.** Arcualia: (0) absent, (1) present. Ch.61 of Keating & Donoghue (2016).
**215.** Transversely biting teeth: (0) absent, (1) present. Ch.64 of Keating & Donoghue (2016).
**216.** Braincase with lateral walls: (0) absent, (1) present. Ch.67 of Keating & Donoghue (2016).
**217.** Neurocranium entirely closed dorsally and covering the brain: (0) absent, (1) present. Ch.68 of Keating & Donoghue (2016).
**218.** Occiput enclosing vagus and glossopharyngeal nerves: (0) absent, (1) present. Ch.69 of Keating & Donoghue (2016).
**219.** Annular cartilage: (0) absent, (1) present. Ch.70 of Keating & Donoghue (2016).
**220.** Trunk dermal skeleton: (0) absent, (1) present. Ch.73 of Keating & Donoghue (2016).
**221.** Calcified cartilage: (0) absent, (1) present. Ch.75 of Keating & Donoghue (2016).
**222.** Cartilage composed of huge clumped chondrocytes: (0) absent, (1) present. Ch.76 of Keating & Donoghue (2016).
**223.** Superficial (often ornamented) layer of the dermal skeleton: (0) absent, (1) present. Ch.78 of Keating & Donoghue (2016).
**224.** Isopedine in dermal skeleton: (0) absent, (1) present. Ch.81 of Keating & Donoghue (2016).
**225.** Tubular dentine in dermal skeleton: (0) absent, (1) present. Ch.84 of Keating & Donoghue (2016).
**226.** Costulate tubercles: (0) absent, (1) present. Ch.89 of Keating & Donoghue (2016).
**227.** Oak-leaf shaped tubercles: (0) absent, (1) present. Ch.90 of Keating & Donoghue (2016).
**228.** Oral plates: (0) absent, (1) present. Ch.91 of Keating & Donoghue (2016).
**229.** Denticles in pharynx: (0) absent, (1) present. Ch.92 of Keating & Donoghue (2016).
**230.** Dermal head covering in adult state: (0) absent, (1) present. Ch.93 of Keating & Donoghue (2016).
**231.** Massive endoskeletal head shield covering the gills dorsally: (0) absent, (1) present. Ch.96 of Keating & Donoghue (2016).
**232.** Sclerotic ossicles: (0) absent, (1) present. Ch.97 of Keating & Donoghue (2016).
**233.** Ossified endoskeletal sclera encapsulating the eye: (0) absent, (1) present. Ch.98 of Keating & Donoghue (2016).
**234.** High blood pressure: (0) absent, (1) present. Ch.99 of Keating & Donoghue (2016).
**235.** Hyperosmoregulation: (0) absent, (1) present. Ch.100 of Keating & Donoghue (2016).
**236.** Forward migration of post-otic myomeres: (0) absent, (1) present. Ch.102 of Keating & Donoghue (2016).
**237.** Larval phase: (0) absent, (1) present. Ch.103 of Keating & Donoghue (2016).
**238.** Pineal opening: (0) covered, (1) uncovered. Ch.104 of Keating & Donoghue (2016).
**239.** Odontodes: (0) Polyodontode, (1) monodontodes. Ch.112 of Keating & Donoghue (2016).
**240.** Triradiate postbranchial spine: (0) absent, (1) present. Ch.116 of Keating & Donoghue (2016).
**241.** Median dorsal ridge scales: (0) absent, (1) present. Ch.117 of Keating & Donoghue (2016).
**242.** Hook shaped median dorsal ridge scales: (0) absent, (1) present. Ch.118 of Keating & Donoghue (2016).
**243.** Body scales with visceral ribs: (0) absent, (1) present. Ch.120 of Keating & Donoghue (2016). *Continuous Characters*. Continuous characters are normalised so the range of character states is between 0 and 1.
**244.** Ratio of pineal plate width to pineal plate length. Ch.66 of Randle & Sansom (2016) and ch.106 of Randle & Sansom (2017).
**245.** Ratio of pineal plate width to orbital plate median lamellae length. Ch.67 of Randle & Sansom (2016) and ch.107 of Randle & Sansom (2017).
**246.** Ratio of rostral plate width to rostral plate length. Ch.68 of Randle & Sansom (2016) and ch.108 of Randle & Sansom (2017).
**247.** Ratio of rostral plate length to dorsal plate length. Ch.69 of Randle & Sansom (2016) and ch.109 of Randle & Sansom (2017).
**248.** Ratio of the median process of the orbital plate length to distance from orbital opening to orbital opening. Ch.70 of Randle & Sansom (2016) and ch.110 of Randle & Sansom (2017).
**249.** Ratio of posterior process of orbital plate length to orbital plate length. Ch.71 of Randle & Sansom (2016) and ch.111 of Randle & Sansom (2017).
**250.** Ratio of anterior process of orbital plates length to orbital plate length. Ch.72 of Randle & Sansom (2016) and ch.112 of Randle & Sansom (2017).
**251.** Ratio of orbital plate median lamellae length to orbital plate length. Ch.73 of Randle & Sansom (2016) and ch.113 of Randle & Sansom (2017).
**252.** Ratio of orbital plate length to dorsal plate length. Ch.74 of Randle & Sansom (2016) and ch.114 of Randle & Sansom (2017).
**253.** Ratio of the distance of the branchial opening from the anterior end of the dorsal plate to dorsal plate length. Ch.75 of Randle & Sansom (2016) and ch.115 of Randle & Sansom (2017).
**254.** Ratio of branchial plate length to dorsal plate length. Ch.76 of Randle & Sansom (2016) and ch.115 of Randle & Sansom (2017).
**255.** Ratio of cornual plate length to dorsal plate length. Ch.77 of Randle & Sansom (2016) and ch.117 of Randle & Sansom (2017).
**256.** Ratio of dorsal plate width to dorsal plate length. Ch.16 of Pernègre & Goujet (2007), ch.32 Pernègre & Elliott (2008) ch.78 of Randle & Sansom (2016) and ch.118 of Randle & Sansom (2017).
**257.** Ratio of dorsal shield width to dorsal shield length (excluding cornual plates). Ch.79 of Randle & Sansom (2016) and ch.119 of Randle & Sansom (2017).
**258.** Ratio of distance to widest part of the dorsal plate from the anterior end of dorsal plate to dorsal plate length. Ch.80 of Randle & Sansom (2016) and ch.120 of Randle & Sansom (2017).
**259.** Ratio of distance to beginning of embayment (in dorsal plate) area from anterior end of dorsal plate to length of dorsal plate. Ch.81 of Randle & Sansom (2016) and ch.121 of Randle & Sansom (2017).
**260.** Ratio of distance to narrowest part of embayment (in dorsal plate) from the anterior end of the dorsal plate to dorsal plate length. Ch.82 of Randle & Sansom (2016) and ch.122 of Randle & Sansom (2017).
**261.** Ratio of narrowest part of embayment (in dorsal plate) width to dorsal plate width. Ch.83 of Randle & Sansom (2016) and ch.123 of Randle & Sansom (2017).
**262.** Ratio of pineal notch depth in dorsal plate to dorsal plate length. Ch.84 of Randle & Sansom (2016) and ch.124 of Randle & Sansom (2017).
**263.** Ratio of dorsal spine base width to dorsal spine base length. Ch.85 of Randle & Sansom (2016) and ch.125 of Randle & Sansom (2017).
**264.** Ratio of dorsal spine base length to dorsal plate length. Ch.86 of Randle & Sansom (2016) and ch.126 of Randle & Sansom (2017).
**265.** Ratio of dorsal shield width to dorsal shield length. For Cyathaspididae or taxa with a singular dorsal shield. Ch.127 of Randle & Sansom (2017).
**266.** Ratio of orbital width to dorsal shield length. For Cyathaspididae or taxa with a singular dorsal shield. Ch. 128 of Randle & Sansom (2017).
**267.** Ratio of pineal macula-rostral length (from anterior end of the dorsal shield) to dorsal shield length. For Cyathaspididae or taxa with a singular dorsal shield. **Ch.**129 of Randle & Sansom (2017).
**268.** Ratio of orbital length-rostral (from anterior end of dorsal shield) to dorsal shield length. For Cyathaspididae or taxa with a singular dorsal shield. Ch.130 of Randle & Sansom (2017). *Discretised characters.* These are continuous characters that have undergone discretisation (see methods above). All are ordered.
**269.** Ratio of pineal plate width to pineal plate length (discretized character 106), (0) 0.83-3.20, (1) 3.85, (2) 4.33, (3) 4.79, (4) 5.50, (5) 6.00. [Ordered]. Ch.132 of Randle & Sansom (2017).
**270.** Ratio of pineal plate width to orbital plate median lamellae length (discretized character 107), (0) 0.29-1.55, (1) 2.33-2.41. Ordered. Ch.133 of Randle & Sansom (2017).
**271.** Ratio of rostral plate width to rostral plate length (discretized character 108), (0) 0.20, (1) 0.50, (2) 0.64-0.79, (3) 0.94-1.29, (4) 1.42-2.00, (5) 2.14-2.35, (6) 2.54. [Ordered]. Ch.134 of Randle & Sansom (2017).
**272.** Ratio of rostral plate length to dorsal plate length (discretized character 109), (0) 0.10-0.56, (1) 0.70, (2) 0.96. [Ordered]. Ch.135 of Randle & Sansom (2017).
**273.** Ratio of the median process of the orbital plate length to distance from orbital opening to orbital opening (discretized character 110), (0) 0.10, (1) 0.15, (2) 0.20-0.21, (3) 0.23-0.27, (4) 0.30-0.45. [Ordered]. Ch.136 of Randle & Sansom (2017).
**274.** Ratio of posterior process of orbital plate length to orbital plate length (discretized character 111), (0) 0.25, (1) 0.41-0.91. [Ordered]. Ch.137 of Randle & Sansom (2017).
**275.** Ratio of anterior process of orbital plates length to orbital plate length (discretized character 112), (0) 0.09-0.35, (1) 0.40, (2) 0.52. [Ordered]. Ch.138 of Randle & Sansom (2017).
**276.** Ratio of orbital plate median lamellae length to orbital plate length (discretized character 113), (0) 0.23-1.03, (1) 1.26-1.36, (2) 1.63. [Ordered]. Ch.139 of Randle & Sansom (2017).
**277.** Ratio of orbital plate length to dorsal plate length (discretized character 114), (0) 0.13-0.49, (1) 0.59, (2) 0.62. [Ordered]. Ch.140 of Randle & Sansom (2017).
**278.** Ratio of the distance of the branchial opening from the anterior end of the dorsal plate to dorsal plate length (discretized character 115), (0) 0.29, (1) 0.41-0.65, (2) 0.79, (3) 0.90-0.91. [Ordered]. Ch.141 of Randle & Sansom (2017).
**279.** Ratio of branchial plate length to dorsal plate length (discretized character 116), (0) 0.10, (1) 0.20-0.21, (2) 0.28-0.61, (3) 0.71-0.77, (4) 0.83. [Ordered]. Ch.142 of Randle & Sansom (2017).
**280.** Ratio of cornual plate length to dorsal plate length (discretized character 117), (0) 0.15, (1) 0.25-0.44, (2) 0.52-0.56. [Ordered]. Ch.143 of Randle & Sansom (2017).
**281.** Ratio of dorsal plate width to dorsal plate length (discretized character 118), (0) 0.34, (1) 0.45-0.96, (2) 1.05, (3) 1.12, (4) 1.23. Ch.16 of Pernègre & Goujet (2007) and ch.32 Pernègre & Elliott (2008). [Ordered]. Ch.144 of Randle & Sansom (2017).
**282.** Ratio of dorsal shield width to dorsal shield length (excluding cornual plates) (discretized character 119), (0) 0.27, (1) 0.35, (2) 0.42-0.85, (3) 0.93, (4) 1.04-1.06. [Ordered]. Ch.145 of Randle & Sansom (2017).
**283.** Ratio of distance to widest part of the dorsal plate from the anterior end of dorsal plate to dorsal plate length (discretized character 120), (0) 0.26-0.61, (1) 0.68, (2) 0.78, (3) 0.92. [Ordered]. Ch.146 of Randle & Sansom (2017).
**284.** Ratio of distance to beginning of embayment (in dorsal plate) area from anterior end of dorsal plate to length of dorsal plate (discretized character 121), (0) 0.51-0.59, (1) 0.63-0.65, (2) 0.71-0.74. [Ordered]. Ch.147 of Randle & Sansom (2017).
**285.** Ratio of distance to narrowest part of embayment (in dorsal plate) from the anterior end of the dorsal plate to dorsal plate length (discretized character 122), (0) 0.53, (1) 0.64-0.79, (2) 0.93-0.97. [Ordered]. Ch.148 of Randle & Sansom (2017).
**286.** Ratio of distance to narrowest part of embayment (in dorsal plate) from the anterior end of the dorsal plate to dorsal plate length (discretized character 123), (0) 0.51, (1) 0.70-0.91. [Ordered]. Ch.149 of Randle & Sansom (2017).
**287.** Ratio of pineal notch depth in dorsal plate to dorsal plate length (discretized character 124), (0) 0.02-0.08, (1) 0.11. [Ordered]. Ch.150 of Randle & Sansom (2017).
**288.** Ratio of dorsal spine base width to dorsal spine base length (discretized character 125), (0) 0.13-0.46, (1) 0.8. [Ordered]. Ch.151 of Randle & Sansom (2017).
**289.** Ratio of dorsal spine base length to dorsal plate length (discretized character 126), (0) 0.08-0.09, (1) 0.11, (2) 0.14-0.25, (3) 0.28-0.30, (4) 0.32-0.34, (5) 0.36-0.37, (6) 0.4. [Ordered]. Ch.152 of Randle & Sansom (2017).
**290.** Ratio of dorsal shield width to dorsal shield length (discretized character 127), (0) 0.37, (1) 0.46-0.79, (2) 0.87. For Cyathaspididae or taxa with a singular dorsal shield. [Ordered]. Ch.153 of Randle & Sansom (2017).
**291.** Ratio of orbital width to dorsal shield length (discretized character 128), (0) 0.24, (1) 0.28-0.30, (2) 0.34-0.36, (3) 0.40-0.53, (4) 0.56, (5) 0.64. For Cyathaspididae or taxa with a singular dorsal shield. [Ordered]. Ch.154 of Randle & Sansom (2017).
**292.** Ratio of pineal macula length (from anterior end of the dorsal shield) to dorsal shield length (discretized character 129), (0) 0.16-0.18, (1) 0.21-0.30, (2) 0.32. For Cyathaspididae or taxa with a singular dorsal shield. [Ordered]. Ch.155 of Randle & Sansom (2017).
**293.** Ratio of orbital length (from anterior end of dorsal shield) to dorsal shield length (discretized character 130), (0) 0.05, (1) 0.10-0.17, (2) 0.19-0.20, (3) 0.23-0.25. For Cyathaspididae or taxa with a singular dorsal shield. [Ordered]. Ch.156 of Randle & Sansom (2017).

### Phylogenetic Analyses

Parsimony analyses were conducted in TNT using the tree-searching with new technologies and branch breaking (xmult and bb functions see scripts) (Goloboff, Farris, & Nixon, 2008). Bayesian analyses were performed using MrBayes (Ronquist & Huelsenbeck, 2003) (settings in supplementary information). To visualize the large distribution of most parsimonious trees, the Tree Set Viz package (Hillis, Heath, & St. John, 2005) was applied in Mesquite (Maddison & Maddison, 2011). This used pairwise comparisons (rooted Robinson-Fould’s distances) and multidimensional scaling to reconstruct axes of variation for trees.

### Stratigraphic Congruence

The stratigraphic congruence of time-scaled phylogenies was tested using the R ‘strap’ package (Bell & Lloyd, 2015). Non-heterostracan taxa were pruned from the trees in order to assess the performance of heterostracan ingroup taxa. Stratigraphic congruence indices use the temporal distribution of taxa to assess the fit of tree topologies to their constituent taxas’ fossil record (Sansom et al., 2018). Genus level stratigraphic occurrence data was collected from the literature (Sansom et al., 2014, Sallan et al/, 2018). Species level stratigraphic data was included for those taxa with multiple species included in the phylogenetic analysis.

## 6. RESULTS

Initial parsimony searches using all taxa recovered little strict consensus. *Eriptychius* was found to be a wild-card taxon and therefore eliminated for subsequent searches. Furthermore, many psammosteid taxa were taxonomically equivalent which prohibitively increased the number of most parsimonious trees without adding phylogenetic signal. Selected psammosteid taxa were therefore removed from subsequent searches and replaced with an equivalent composite taxon.

Heuristic searches without *Eriptychius* and taxonomically equivalent psammosteid taxa achieved a better resolution but still resulted in high number of most parsimonious trees (Fig. 5a). The Heterostraci and non-heterostracan Pteraspidimorphi (*Arandaspis, Sacabambaspis* and *Astraspis*) do not form a monophyletic clade in the strict consensus and form a polytomy as the sister group to non-Pteraspidimorphi gnathostomes (total group). The Heterostracans were recovered as monophyletic in all instances.

**Figure 5.**
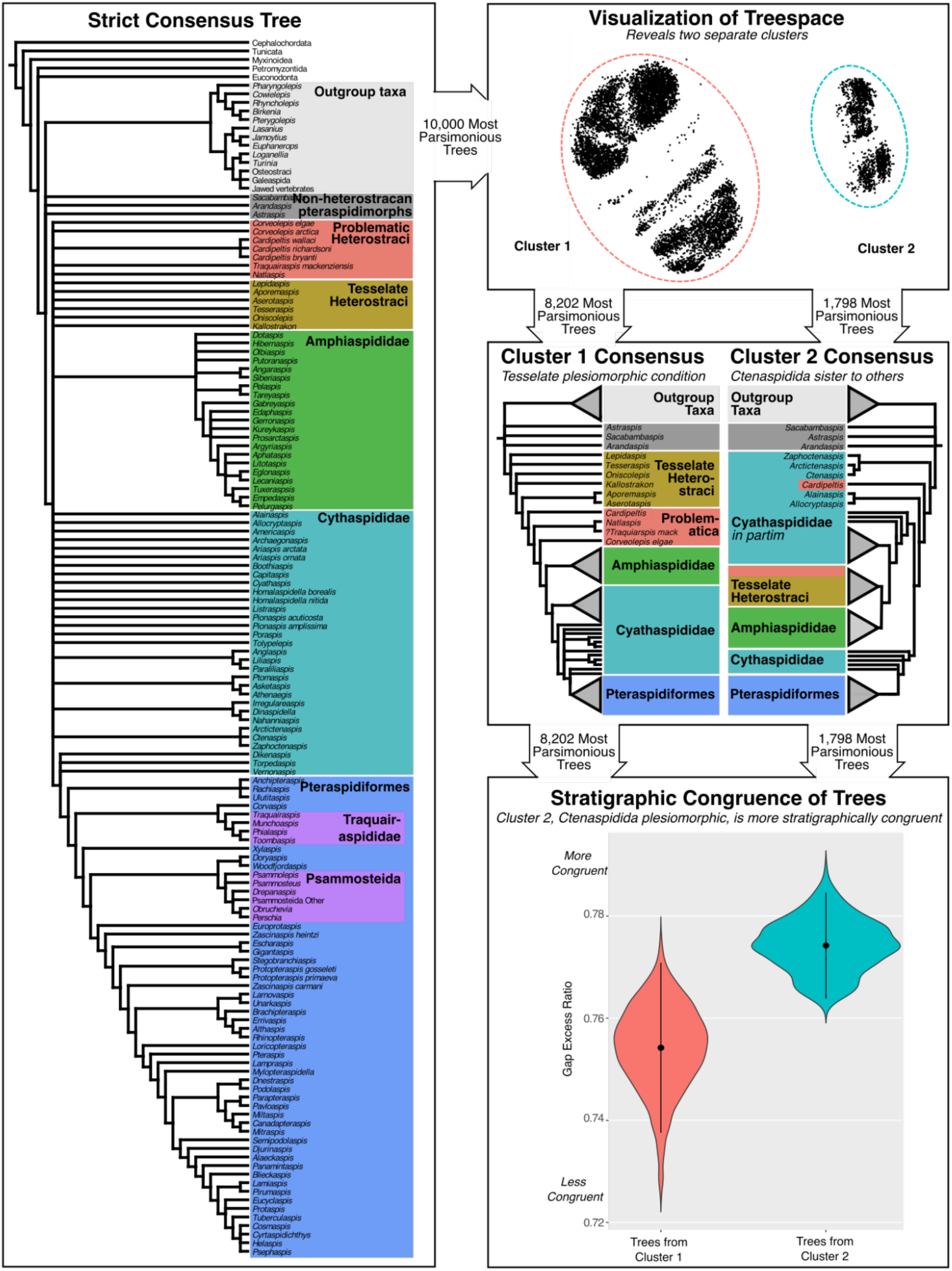
Results from the parsimony phylogenetic analyses. A. Strict consensus tree of 10,000+ most parsimonious trees. B. Tree visualization of 10,000 most parsimonious trees (Multi-dimensional scaling using Robinson-Fould’s distances, C. Simplified consensus trees showing the relationships within each of the two clusters, D. The stratigraphic congruence of most parsimonious trees from each cluster evaluated using the Gap Excess Ratio.

Characters supporting the position of heterostracans on the gnathostome stem include the presence of: a calcified dermal skeleton, olfactory peduncles, otic capsule anterior to branchials, vertical semicircular canals forming loops that are separate from the roof of the utriculus, endodermal gill lamellae and a trunk dermal skeleton. Synapomorphies supporting a monophyletic Heterostraci include: the presence of the supra-orbital commissure (CM.SO) canal joining the dorsal lateral canal to the inter-orbital or supra-orbital (SOC) canal and a single pair of branchial openings.

The strict consensus tree recovers monophyly of some important groups i.e. Amphiaspididae and Pteraspidiformes (Fig. 5A). The Psammosteida and Traquairasididae are also monophyletic, both as part of the Pteraspidiformes. A lack of resolution exists in the relationships within and between the other major groups i.e. Cyathaspididae, tesselate Heterostraci and the problematic taxa. To evaluate these relationships and identify underlying topologies, we applied tree space visualization using the ‘Tree Set Viz’ module in Mesquite (Maddison and Maddison 2004) to 10,000 most parsimonious trees obtained using the discretised quantitative characters. Two separate clusters of trees were identified (Fig. 5b). The main difference between the clusters is the relative rooting solution of the Heterostraci. In one cluster (Fig. 5c, n=8,202), the tessellate heterostracans (*Lepidaspis, Tesseraspis, Oniscolepis, Kallostrakon, Aporemaspis* and *Aserotaspis*) form a paraphyletic sister-group to all other Heterostraci, whilst the Cyathaspididae, Amphiaspididae and Pteraspidifromes form a large clade. In the second cluster (n=1,798), the ctenaspids (*Ctenaspis, Arctictenaspis* and *Zaphoctenaspis*) are sister group to all other heterostracans and the tesselate forms are nested. In both solutions the Cyathaspididae are paraphyletic to the Pteraspidiformes, but in cluster 2, they are also paraphyletic with respect to the tesselate forms and Amphiaspididae (Supplementary Figure 1).

To distinguish between these two equally parsimonious clusters with very different rooting solutions for Heterostraci, we applied stratigraphic congruence estimates. All topologies resulting from the different coding methods have a better fit to stratigraphy than those generated from randomly permutated trees, but trees from cluster 2 have the best fit to stratigraphy in all instances i.e. the Ctenaspididae as sister group to all other heterostracans (Fig.5d).

The Bayesian analysis yields a very unconventional result for heterostracan intra- and inter-relationships as well as vertebrate phylogeny: the heterostracans are recovered as paraphyletic stem-vertebrates. The tessellate heteostracans and Ordovician pteraspidimorphs are successive sister groups to all non-heterostracan vertebrates. The majority rule consensus tree (supplementary Fig. 2) fails to recover any relationships within the Heterostraci, with many heterostracan taxa in a large polytomy.

## 7. DISCUSSION

Results from parsimony phylogenetic analyses place the Heterostraci and the Ordovician Pteraspidimorphi as the deepest branching skeletonizing vertebrates. This is in accordance with the results of Donoghue *et al*. (2000) and Keating & Donoghue (2016). This is contra to pteraspidimorph positions in other phylogenetic analyses, for example as sister group to galeaspids + osteostracans + jawed vertebrates in Sansom *et al*. (2010) and Gess *et al*. (2006), and as the sister group to thelodonts + galeaspids + osteostracans + jawed vertebrates in Blom (2012). Within the Pteraspidimorphi, our greater taxon sampling and different character models are unable to support either of the contested Ordovician taxa as sister group to Heterostraci i.e. Arandaspida or Astraspis (Donoghue *et al*. 2000; Sansom, 2009; Sansom *et al*., 2010; Gabbott *et al*., 2016; Keating & Donoghue, 2016). Instead our results are unresolved at this node (similar to Gess et al. 2006). Further examination of these incomplete and seemingly conflicting taxa may help to resolve this conflict.

Within the Heterostraci, it first appears that there is conflict and little resolution of relationships, but this is resolved through the application of tree space visualization. Relationships within the Heterostraci are actually relatively stable, but masked because there are two equally parsimonious solutions for the rooting of the group (fig. 5). Of those two solutions, we find the Ctenaspididae as sister group to all other heterostracans to be the preferred solution (fig. 6) because this topology is much more congruent with the stratigraphic distribution of taxa, relative to the more traditional solution i.e. the tesselate/micromeric condition being plesiomorphic with *Tesseraspis* as sister taxon to all other heterostracans (Halstead 1973). This is primarily a result of the position of *Athenaegis* (one of the stratigraphically oldest heterostracans arising in the Wenlock) and other Silurian Cyathaspididae being positioned in a derived position with regards to the Devonian Amphiaspididae (Lochkovian-Pragian) in cluster 1, whereas, this is the other way round in cluster 2.

**Figure 6.**
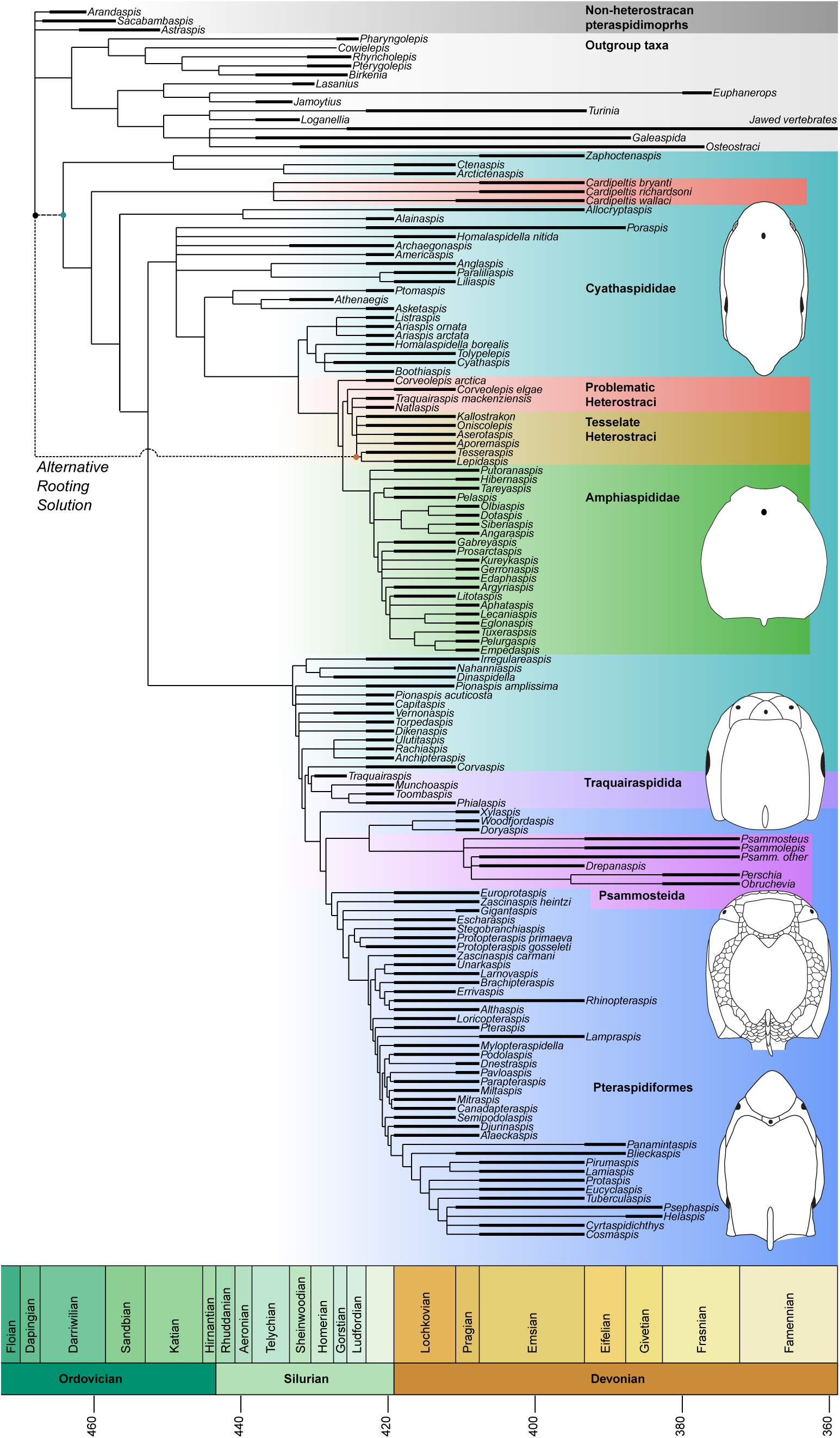
Phylogeny of Heterostraci, time scaled against stratigraphic age range data. The ‘Ctenaspididae basal’ rooting solution is shown here, with the alternative ‘tesselate basal’ rooting solution highlighted

Of the traditional Heterostracan clades, the Amphiaspididae are monophyletic in all analyses. In instances when the Ctenaspididae are sister group (cluster 2) to all other heterostracans (fig. 5, Fig. 6), the Amphiaspididae are sister group to the tessellate and problematic heterostracans rather than the Cyathaspididae as envisioned by Obruchev (1967), Halstead (1973), Novitskaya (1983), Blieck *et al*. (1991) and Janvier (1996). However, the Amphiaspididae and problematica are in a much larger group also containing many members traditionally assigned to the Cyathaspididae. In the tessellate-basal model (cluster 1), the Amphiaspididae are the next successive sister group. Therefore, a close affinity (sister group) between the Amphiaspididae and Cyathaspididae is rejected here.

The placement of Psammosteida within the Pteraspidiformes is only retained, *sensu stricto,* if the Anchipteraspididae are retained as the deepest branching Pteraspidiformes. This pattern is consistent with the phylogenetic positioning as demonstrated by Blieck (1984), Pernègre & Goujet (2007) and Randle & Sansom (2017). This analysis did not provide much insight into the inter-relationships or monophyly of the Psammosteidae, but this has been resolved in thorough detail by Glinkskiy (2017).

In parsimony analyses, the Traquairaspididae *sensu* Tarrant (1991) is retained, with the exception of *Rimaventeraspis* in the Ctenaspididae-basal model trees. The close association of Traquairaspididae, Psammosteidae and the Pteraspidiformes recovered in our phylogeny was hypothesized by Halstead (1973). The Cyathaspididae *sensu* Denison (1964) were found to be polyphyletic.

A monophyletic Cyathaspididae has been hypothesized by many authors including Halstead (1973), Blieck *et al*. (1991) and Janvier (1996); a phylogeny by Lundgren & Blom (2013) only including cyathaspid taxa recovered similar relationships to those of Denison (1964). However, recent phylogenetic analyses including pteraspid and cyathaspid taxa demonstrated the Cyathaspididae to be paraphyletic (Randle & Sansom, 2017). Based on similarity of headshields, the Cyathaspididae and Amphiaspididae were believed to have a close affinity (Obruchev, 1967; Halstead, 1973; Novitskaya, 1983, 2004, 2008; Blieck *et al*., 1991; Janvier, 1996). This solution is not recovered in any of our analyses. Instead the Cyathaspididae are paraphyletic with both heterostracan rooting solutions (fig. 5). The relationships of Pteraspidiformes recovered from the parsimony analyses are the same as those recovered in Randle & Sansom (2017). The Anchipteraspididae are not recovered here as sister group to the pteraspids but rather as the sister group to the multi-plated forms i.e. Psammosteidae + Traquairaspididae + pteraspids.

The position of heterostracans with previously uncertain affinities produces fairly consistent patterns. *Cardipeltis*, in all instances, is positioned as the sister group to the clade containing Cyathaspididae, Pteraspidiformes, Traquairaspididae, Amphiaspididae and Psammosteidae. The Corvaspididae *sensu* Loeffler & Dineley (1976), which included the Canadian Arctic *Corveolepis arctica* and *Corvaspis*, is not recovered as monophyletic as suggested by Blieck & Karatajūtė-Talimaa (2001). Although no articulated specimens of *Corvaspis* have ever been found, this relationship is unlikely to change with the discovery of an articulated specimen. *Corvaspis* is composed of separate plates, whereas *Corveolepis* has a dorsal headshield composed of a singular plate. It is clear that *?Traquairaspis mackenziensis* does not have close affinity to the monophyletic Traquairaspididae. Our results indicate *?Traquairaspis* has a close affinity with *Natlaspis*, another problematic heterostracan from the Canadian Arctic: both taxa have their orbits and branchial openings enclosed by the dorsal headshield and an uncovered pineal organ. Dineley & Loeffler (1976) placed *?Traquairaspis* in the Traquairaspididae due to similarities in their external ornament, however, it is most likely that these Canadian traquairaspids represent a new and as yet un-described group of heterostracans.

Creating a phylogeny including disparate clades needs an identification or interpretation of homologous features. This can be difficult for fossil taxa for which we may have no embryological and developmental data or any morphological intermediates. Therefore, it’s up to us to make judgments and interpretations of fossil taxas’ anatomy. A distinction made here was between taxa having a dorsal headshield and a dorsal plate. The presence of a dorsal headshield was coded if the orbital and pineal openings, along with the rostral areas were encompassed within one plate. Whereas, if these features were delimited by different plates, then taxa were coded as having a dorsal plate. Many subsequent characters were contingent upon those two features. However, not enough is known about the growth and development of heterostracan fishes dermal skeleton, to say if the dorsal plate of Traquairaspididae is homologous to that of Pteraspidiformes and Psammosteidae.

Results obtained from the Bayesian phylogenetic analyses are unusual and have not been recovered in any other phylogenetic analyses to date (although the majority of these are parsimony based) (Donoghue *et al*., 2000; Gess *et al*., 2006; Blom, 2012; Gabbott *et al*., 2016; Keating & Donoghue, 2016). The Bayesian trees recover the Heterostraci as paraphyletic and as stem-vertebrates, which would have great implications for character evolution in this part of the vertebrate evolutionary tree. Due to this unconventional topology, along with the placement of conodonts, cyclostomes and Ordovician pteraspidimorphs in a more derived position within the vertebrate tree, this topology is treated with caution. Greater taxon inclusion of other stem-gnathostome clades may produce phylogenies more in line with parsimony analyses; however, this will only be elucidated by future studies.

Phylogenetic analyses incorporating long extinct taxa often incorporate large amounts of uncertainty, especially regarding anatomical interpretations, missing data and large amounts of homoplasy. Despite this, creating evolutionary hypotheses of relationships are vital to unravelling macroevolutionary patterns of diversity and character evolution. This is true for early vertebrates and the Heterostraci, which we present here. It is hoped that with this framework of heterostracan intra-relationships future investigation into the macroevolutionary patterns of early vertebrates will now be possible. Stratigraphic congruence indicates the *Ctenapsis-*rooted tree has the most consistent fit with the temporal fossil record of heterostracans, however, better preserved tessellate specimens would allow a clearer interpretation of their anatomy and would perhaps provide more conclusive answers.

## 8. CONCLUSIONS

Here we present the first phylogenetic analyses of Heterostraci with a broad range of ingroup and outgroup taxa. This tested the monophyly of many clades within the Heterostraci including the Amphiaspididae, Traquairaspididae, Psammosteidae and Cyathaspididae and their relationships to each other. Other taxa included were the problematic and tessellate heterostracans, for which no formal phylogenetic analysis had been conducted. The inclusion of non-heterostracan vertebrates as outgroup taxa also enabled us to reconstruct the position of the Heterostraci on the gnathostomes stem. Results from parsimony analyses found the Amphiaspididae, Traquairaspididae and Pteraspidiformes to be monophyletic. The multi-plated heterostracans (Pteraspidiformes + Psammosteidae + Traquairaspididae) were also recovered in a clade. The Cyathaspididae in all instances were recovered as polyphyletic, indicating a re-interpretation of their relationships is needed. Conflict surrounds the plesiomorphic condition for the Heterostraci, with two models apparent: the ‘tessellate-basal model’ and the ‘ctenaspid-basal-model’. Stratigraphic congruence indices indicate discretising heterostracan quantitative characters produces the most stratigraphically congruent trees - with the highest values seen in the ‘ctenaspid-basal-model’ topologies. The position of the Heterostraci on the gnathostome stem remains ambiguous as they are placed in a polytomy with the Ordovician Pteraspidimorphi as the deepest branching skeletonising vertebrates. Our results suggest discretising quantitative characters can greatly improve the resolution of phylogenetic trees of early vertebrates. We also recommend exploring tree topologies of seemingly poorly resolved strict consensus trees using tree visualization, as there may be genuine, but conflicting signal amongst the most parsimonious solutions.

## Supplementary Data and Figures

**Supplementary Figure 1.**
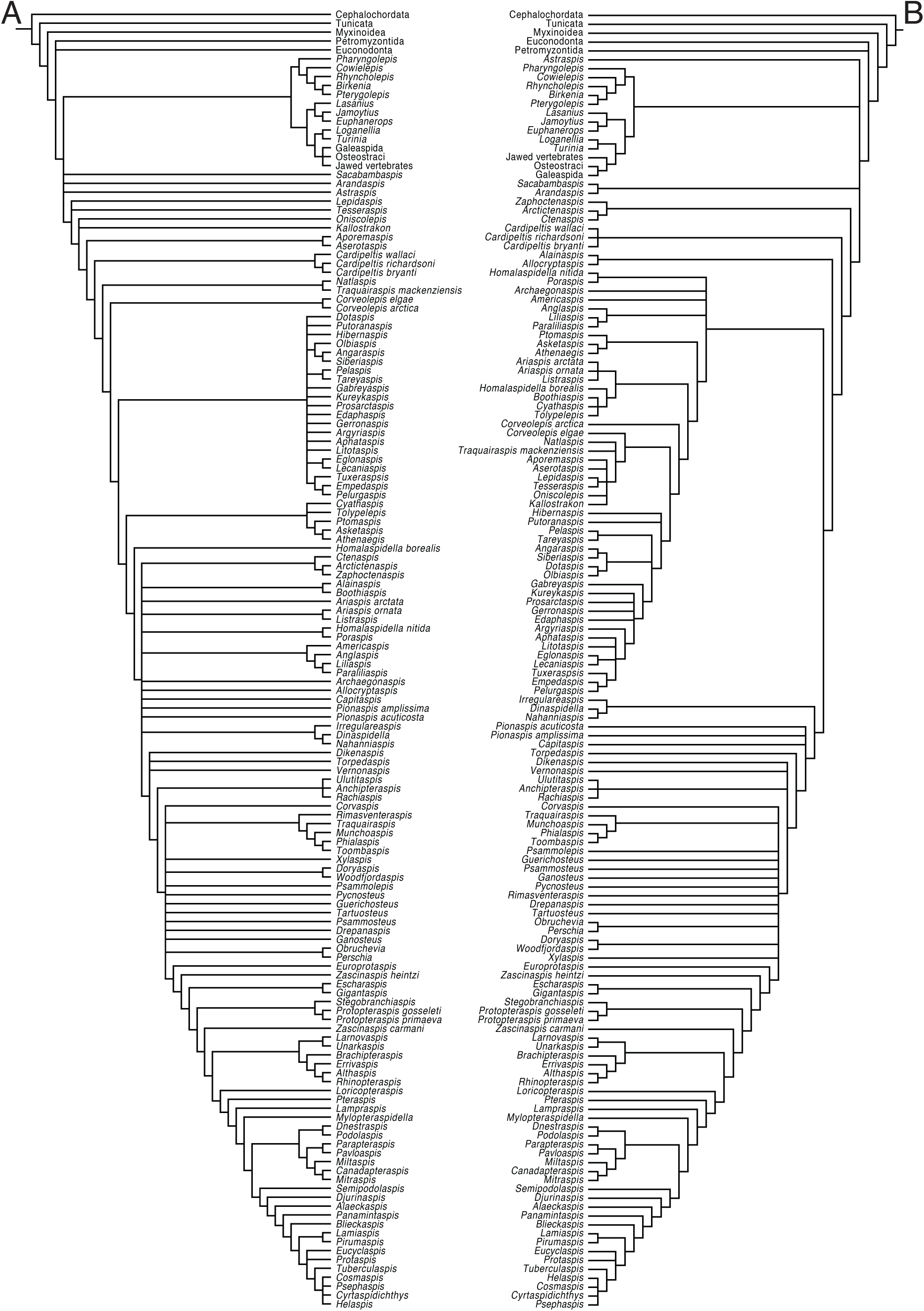
Strict consensus of the two clusters identified using tree visualization: A ‘tesselate basal’ model, B, ‘Ctenaspididae basal model’.

**Supplementary Figure 2.**
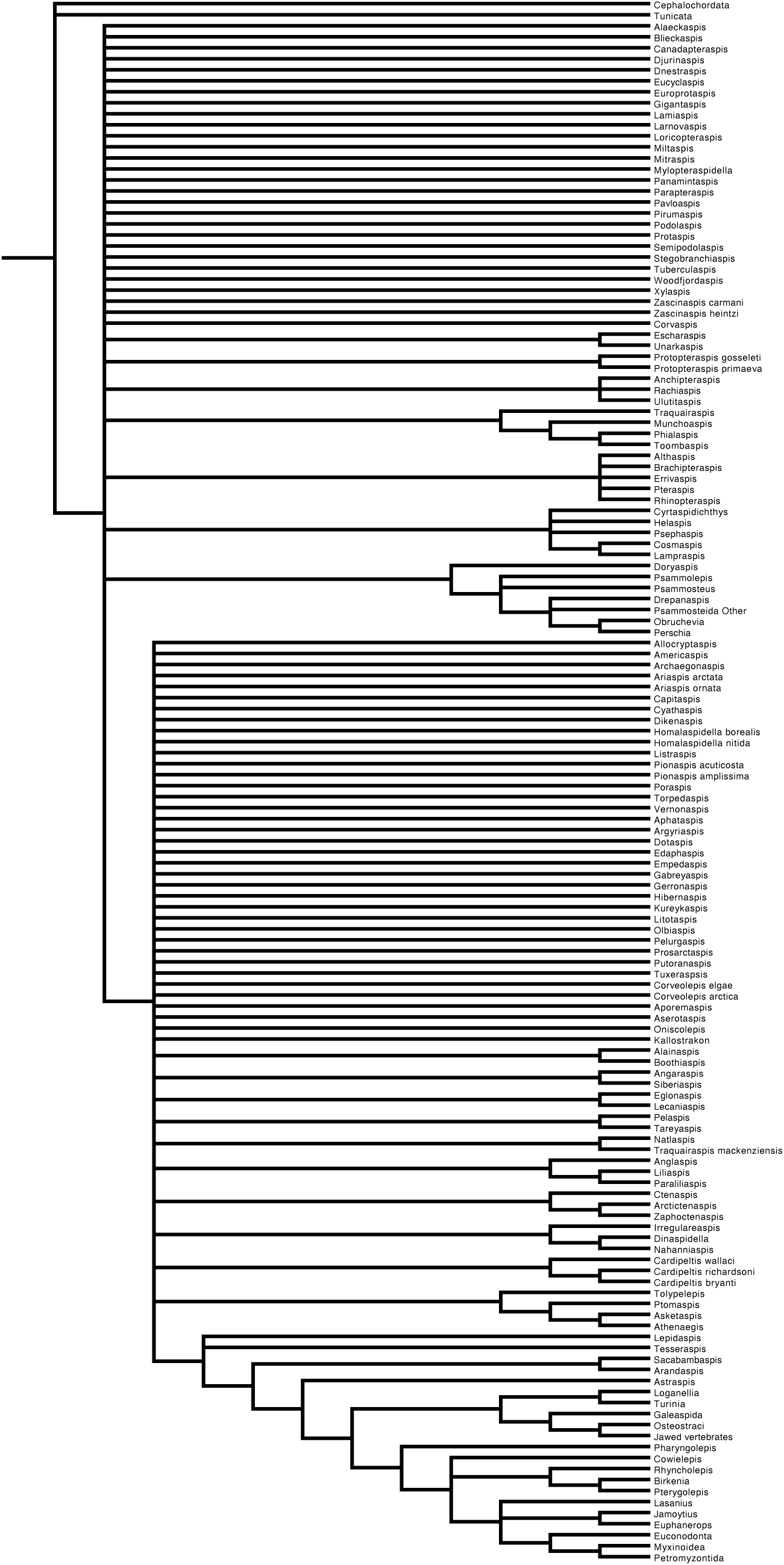
Majority rule consensus tree resulting from Bayesian tree search

Supplementary Data 1. Phylogenetic data matrix in TNT format with TNT search commands

Supplementary Data 2. Phylogenetic data matrix in nexus format with Mr Bayes search commands

## Acknowledgements

This work was completed as part of Emma Randle’s PhD funded by the Natural Environment Research Council (NERC NE/K500859/1) and supported by funding from Synthesis (visit to Naturhistoriska Riksmuseet SE-TAF-3875). We thank the staff at all museums for access to specimens, including Dr Olga Afanassieva and Dr Larissa Novitskaya (Palaeontological Institute, Moscow), Dr Tiiu Märss and Ursula Toom (Geological Institute, Tallinn), Jonathan Clatworthy and Dr Ivan Sansom (Lapworth Museum of Geology, University of Birmingham), Emma Bernard (Natural History Museum, London), William Simpson and Akiko Shinya (Field Museum, Chicago), Kiaeron Sheppard and Margaret Curry (Canadian Museum of Nature) and Dr Lars Werdelin and Jonas Hagström (Naturhistoriska riksmuseet, Stockholm). We also thank Dr Vadim Glinskiy and Tormi Tuuling for the help in acquiring papers. Feedback on earlier version of this work was provided by Russell Garwood and Zerina Johanson. We also thank Peter Tarrant for his support with fieldwork and specimen access.

